# Copy-number dosage regulates telomere maintenance and disease-associated pathways in neuroblastoma

**DOI:** 10.1101/2022.08.16.504100

**Authors:** Martin Burkert, Eric Blanc, Nina Thiessen, Christiane Weber, Joern Toedling, Remo Monti, Victoria M Dombrowe, Maria Stella de Biase, Tom L Kaufmann, Kerstin Haase, Sebastian M Waszak, Angelika Eggert, Dieter Beule, Johannes H Schulte, Uwe Ohler, Roland F Schwarz

**Author notes:** Centre for Molecular Medicine Norway (NCMM), Nordic EMBL Partnership, University of Oslo and Oslo University Hospital, Oslo, Norway.

## Abstract

Telomere maintenance in neuroblastoma is linked to poor outcome and caused by either TERT activation or through alternative lengthening of telomeres (ALT). In contrast to TERT activation, commonly caused by genomic rearrangements or MYCN amplification, ALT is less well understood. Alterations at the ATRX locus are key drivers of ALT but only present in ∼50% of ALT tumors.

To identify potential new pathways to telomere maintenance, we investigate allele-specific gene dosage effects from whole genomes and transcriptomes in 115 primary neuroblastomas. We show that copy-number dosage deregulates telomere maintenance, genomic stability, and neuronal pathways and identify upregulation of variants of histone H3 and H2A as a potential alternative pathway to ALT. We investigate the interplay between *TERT* activation, overexpression and copy-number dosage and reveal loss of imprinting at the *RTL1* gene associated with poor clinical outcome.

These results highlight the importance of gene dosage in key oncogenic mechanisms in neuroblastoma.

## Introduction

Neuroblastoma is the most common extracranial solid tumor in children accounting for 6-10% of malignancies ^1^ and 9% of pediatric cancer deaths ^2^. The disease shows heterogeneous clinical manifestations ranging from high-risk cases with poor survival rates despite multimodal treatment to tumors that spontaneously regress without intervention ^3^. Incidence is highest in the first year of life and only 5% of diagnoses are made in patients older than ten years ^1^. Diagnosis at an advanced age is generally associated with worse outcomes ^2^.

Genetically, neuroblastoma is characterized by low single-nucleotide variants (SNV) burdens and only few recurrently mutated genes ^4^, but frequent somatic copy-number alterations (SCNAs) ^5–7^. Amplification of the oncogenic transcription factor *MYCN*, often through extrachromosomal circular DNAs ^8,9^, is found in 20% of tumors and a key clinical indicator for high-risk disease and poor prognosis ^3,10^. In addition, recurrent segmental gains and losses, including 17q gains and losses of 1p and 11q ^6,11,12^ are associated with unfavorable outcomes ^13^. These SCNAs affect cellular phenotypes by modulating gene expression. Amplifications of *MYCN* and *ALK* upregulate these oncogenes and their downstream targets ^14,15^, and larger segmental gains and losses were also found to correlate well with local RNA levels ^16,17^, which in turn predict patient survival ^14,15,17^.

Telomere maintenance leading to replicative immortality ^18^ is a common mechanism in high-risk neuroblastoma ^19–21^. Canonical telomere maintenance involves activation of the Telomerase reverse transcriptase (*TERT*) gene either indirectly as a downstream effect of *MYCN* amplification, or directly through genomic rearrangements at the *TERT* locus ^19,21^. Alternative lengthening of telomeres (ALT) in tumors that lack *TERT* activation ^22^involves DNA recombination induced by breaks at telomeric sequences ^23^ and is characterized by single stranded telomeric (CCCTAA)_n_ sequences ^24^. Generally, ALT is associated with loss of function mutations in the *ATRX* and *DAXX* genes ^25^and has been found in 50% of all cancer types of the Pan-Cancer Analysis of Whole Genomes (PCAWG) cohort ^26^. Affected tumors show excess telomere length compared to normal tissue and other tumors, including those with activated *TERT* ^26^. In neuroblastoma ALT is associated with ATRX alterations ^27–29^, significantly enriched in relapse cases and associated with poor outcome independent of other risk markers ^20,28^. While previous studies have highlighted the molecular characteristics of telomere maintenance in neuroblastoma ^19,27,28,30,31^, *ATRX* mutations were only found in 25% of high-risk and 50-60% of ALT-positive neuroblastomas ^27,28,32^, suggesting additional yet unrecognized mechanisms of ALT activation. Telomere maintenance is therefore a key phenotypic property of neuroblastoma cells and a prime example of phenotypic convergence in cancer evolution ^33^, where multiple somatic aberrations act individually or in concert to activate telomere maintenance pathways by modulating gene expression.

To reveal such mechanisms, we here investigate the effect of genomic instability on total and allele-specific gene expression and telomere maintenance in 115 primary neuroblastomas. We analyze whole genome sequencing (WGS) and RNA-seq from tumors and WGS of matched normals, characterize local genetic effects on gene expression variability, and examine the role of copy-number dosage in telomere maintenance and survival.

## Results

### Cohort overview

We assembled a cohort of matched tumor WGS and RNA-seq and normal WGS from blood from 115 primary neuroblastoma samples, including 52 samples from the University Hospital of Cologne, previously reported in ^19^, and 63 new specimens from the GPOH-NB2004 clinical trial. All samples were jointly processed using unified pipelines to limit cohort-specific biases (Methods) and stratified according to the GPOH-NB2004 clinical trial protocol ^34^ into 66 high-risk, 6 medium-risk, and 43 low-risk tumours (S.Fig. 1) and equipped with clinical annotations including age, sex and survival times (S.Table 1).

**Figure 1:**
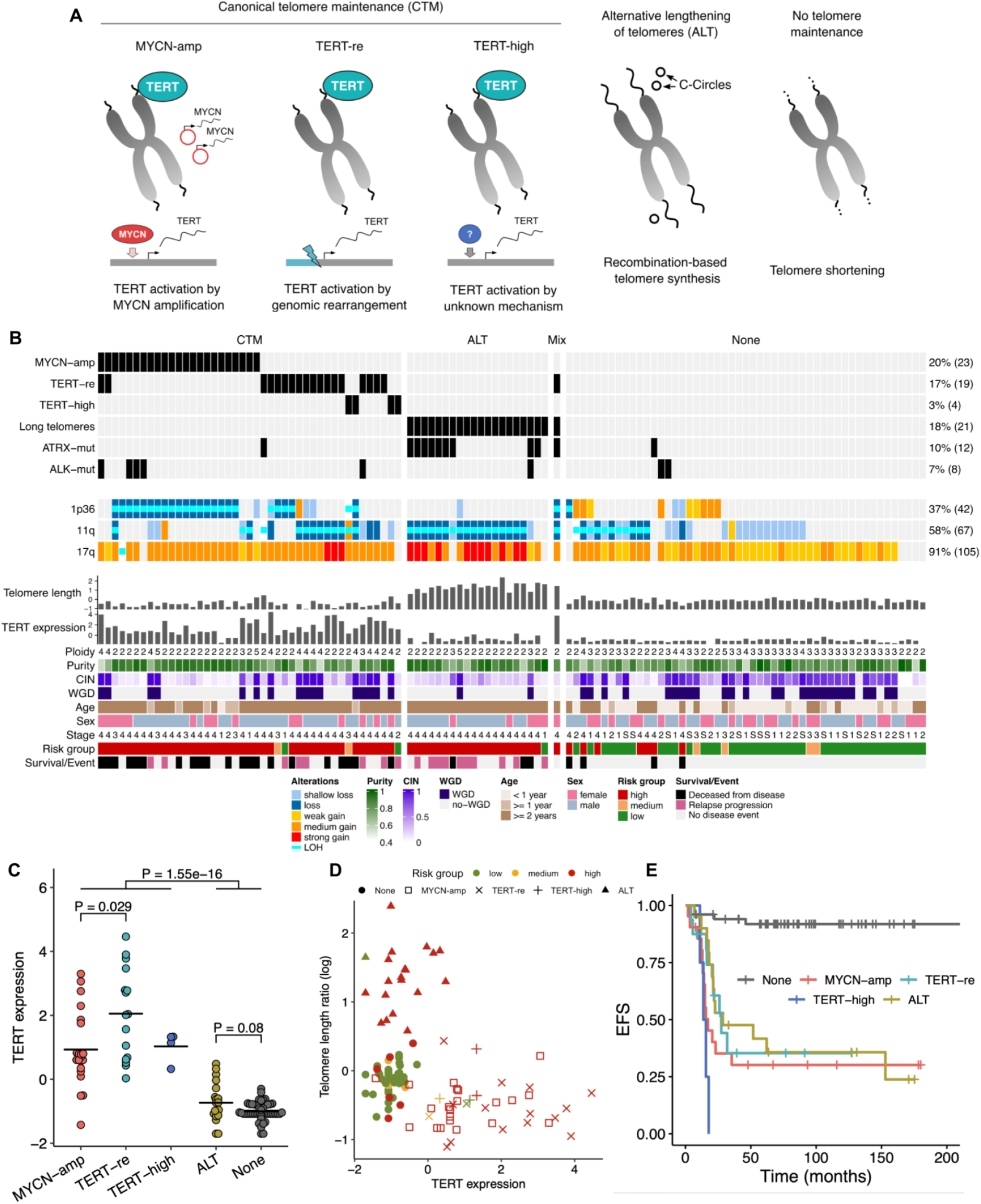
Molecular profiles and telomere maintenance in 115 neuroblastoma tumors. **A** Overview of telomere maintenance mechanisms: In canonical telomere maintenance *TERT* is activated by *MYCN* amplification, genomic-rearrangements at the *TERT* locus or an unknown mechanism. ALT is characterized by excess telomere length and C-circles. Absence of telomere maintenance leads to telomere shortening. **B** Molecular and clinical characterization of 115 neuroblastoma primary tumors (columns). ALT: Alternative lengthening of telomeres, *MYCN*-amp: *MYCN* amplification, *TERT*-re: *TERT* rearrangement, *ATRX*-mut: *ATRX* mutation, *ALK*-mut: *ALK* mutation, LOH: Loss of heterozygosity. WGD: Whole-genome doubling, CIN: Chromosome instability index. **C** Comparison of *TERT* expression between tumors by molecular telomere maintenance characteristics. **D** Telomere length and *TERT* expression per sample. **E** Kaplan-Meier estimate of event free survival (EFS) by telomere maintenance mechanism in primary tumors. Tumors with more than one molecular characteristic are not shown.

Normal samples from blood were genotyped and phased at common germline variant sites (S.Methods). Total and allele-specific gene expression (ASE) was quantified using phased variants and variant effects on gene expression in *cis* were quantified by genome-wide expression quantitative trait locus (eQTL) mapping (S.Methods) ^35^. To explore the mutational landscape we determined somatic single-nucleotide variants (SNVs), structural variants (SVs) and allele-specific somatic copy-number alterations (SCNAs) from WGS (Methods).

### Telomere maintenance status of 115 primary neuroblastomas

We first set out to determine the primary telomere maintenance mechanism (Fig 1A) and genetic alterations across all 115 tumors by examining somatic SNV, SV, SCNA and expression data as well as WGS-based estimates of telomere length (Methods). We found *MYCN* amplifications in 23 tumors (20%), rearrangements affecting the *TERT* locus in 19 tumors (17%) and *ATRX* mutations in 12 tumors (10%), comprising 7 focal deletions, 4 missense or nonsense mutations and one tumor affected by a structural rearrangement (NBL54) (Fig 1B, S.Fig. 2). In addition, *ALK* mutations were found in 8 tumors (7%), of which 6 carried a missense mutation and 2 were affected by genomic amplifications. We queried *TERT* gene expression in all tumors and found both *MYCN* amplified and *TERT*-rearranged samples to have significantly higher *TERT* expression than those lacking both molecular features (Fig. 1C), in line with previous observations ^19,36^. We additionally found 4 tumors without *MYCN* amplification or *TERT* rearrangements to show *TERT* overexpression (S.Fig 5, Methods). To determine the ALT status of tumors we estimated telomere lengths relative to the matched normal tissue by the abundance of telomeric repeat sequences from WGS (S.Fig. 3A, Methods) ^37^. We found 21 tumors to show increased telomere lengths, of which we assigned 20 to the ALT group, as one (NBL54) also harbored a *TERT* rearrangement and upregulation of *TERT* (Fig. 1B, S.Fig 3A,B). We validated our ALT classification by comparison against experimentally determined status of ALT-associated PML-nuclear bodies (APB) ^20^ in 52 donors (S.Fig. 3B) and found a strong correspondence (P = 5.47 × 10^−9^, one-sided Fisher’s Exact Test; sensitivity: 0.86; specificity: 0.97). While *ATRX* altered samples had significantly longer telomeres (P = 1.72 × 10^−6^, one sided Wilcoxon rank sum test) (S.Fig. 4), in 11 out of 20 ALT samples (55%) *ATRX* mutations were not detected, pointing towards alternative activation of the ALT pathway independent of *ATRX* mutations.

**Figure 2:**
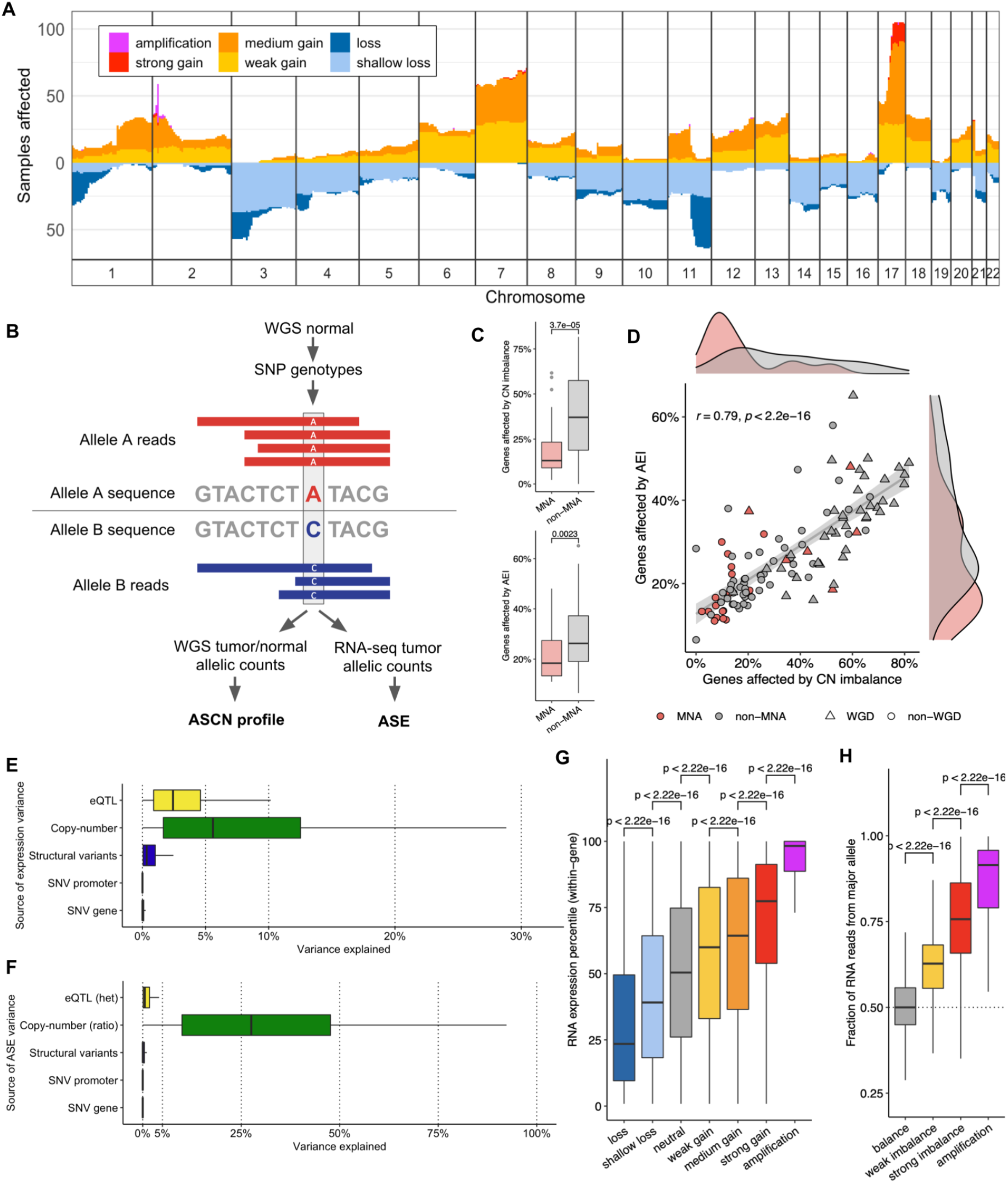
Genetic- and allelic dosage effects in gene regulation. **A** Number of samples affected and copy-number state summarized in 5 Mb genomic bins on chromosomes 1-22. **B** Allele-specific profiling of somatic copy-number and gene expression based on allelic counts at heterozygous SNPs in WGS and RNA-seq. **C** Number of genes affected by copy-number imbalance and AEI for samples with and without *MYCN*-amplification (MNA). Boxplot midlines in (B,C,E,F,H) mark median; upper and lower hinges extend to first and third quartile; upper and lower whiskers extend to the smallest and largest value max. 1.5 × IQR; p-value of two-sided Wilcoxon test is shown between groups. **D** Number of genes affected by copy-number imbalance and allelic expression imbalance (AEI) per sample. CN, copy-number. Quantification of genetic effects on the variance of (**E**) total expression and (**F**) allele-specific expression (ASE). **G** Distribution of within-gene expression percentile per sample and gene by copy-number state. **H** Proportion of RNA reads from major allele per sample and gene by copy-number balance state.

**Figure 3:**
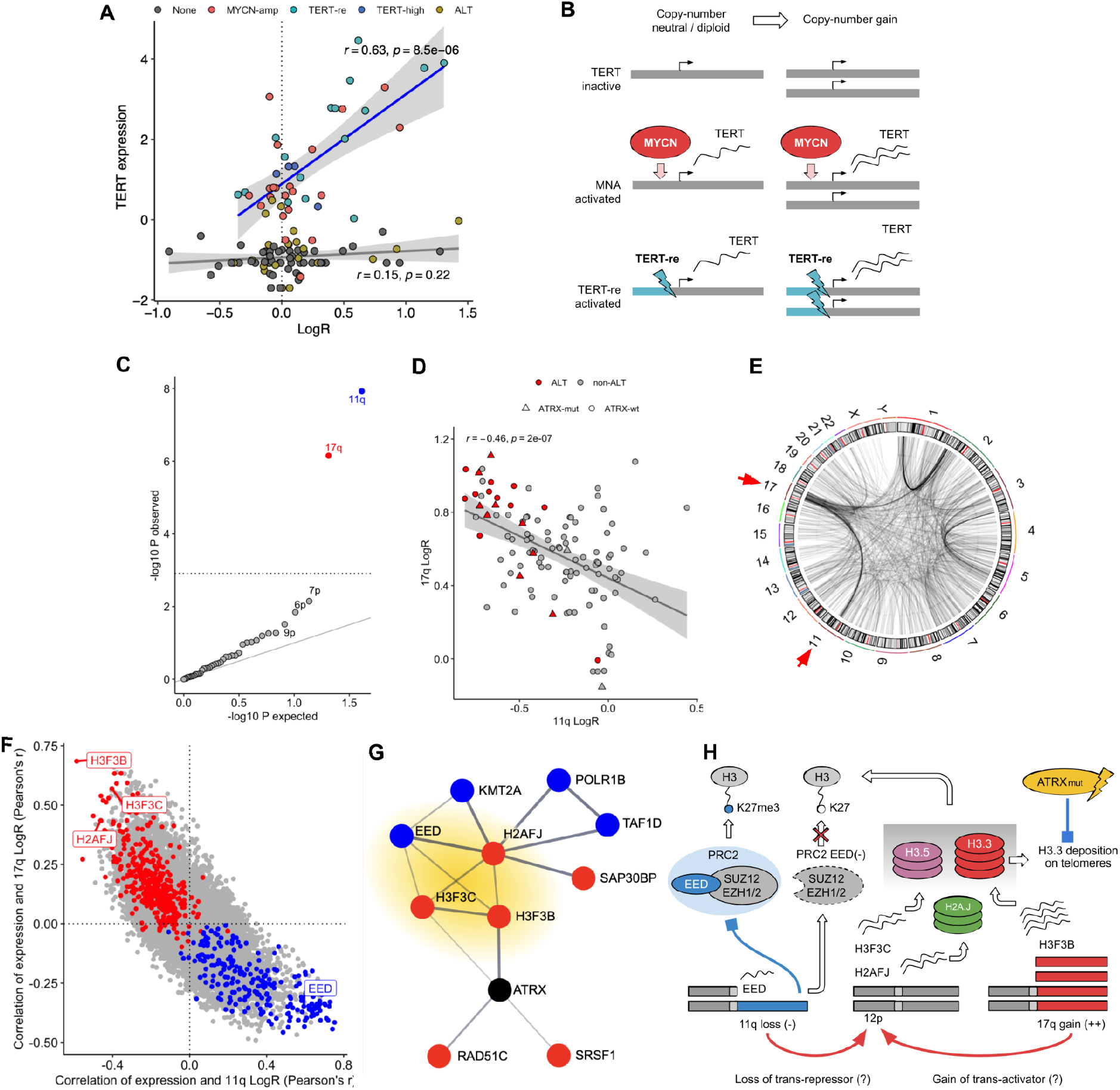
Copy-number dosage effects in telomere maintenance. **A** Copy-number LogR and expression of *TERT* per sample. Regression line of samples with canonical telomere maintenance (*MYCN*-amp, *TERT*-re, *TERT*-high) in blue, regression line of other samples in gray. Gray ribbons indicate 95% confidence intervals. **B** Copy-number gains induce higher *TERT* expression in tumors of activated *TERT*. **C** Association p-values of ALT and coverage LogR per chromosome arm. Significant observations in red (gain) and blue (loss), others in gray. Significance threshold (FWER 0.05) demarcated by gray dotted line. **D** 11q and 17q LogR per sample indicating ALT and *ATRX* status. **E** Genome-wide somatic structural variant breakpoints. Breakpoints of frequent rearrangements between chromosome arms 11q and 17q highlighted by red arrows. **F** Correlation of gene expression and LogR of 17q and 11q. Differentially expressed genes in red (ALT up) and blue (ALT down). Three histone variant genes are upregulated in ALT and substantially correlate with 11q and 17q LogR. **G** STRING database protein interaction network of *ATRX, H3F3B, H2AFJ* and *H3F3B* and their first order interactions among differentially expressed genes (ALT). Red and blue indicate up- and downregulated genes respectively. *ATRX* is not differentially expressed (black). Histone variant genes and *EED* highlighted in yellow. **H** Proposed model of chromatin factor deregulation in ALT: 11q and 17q copy-number alterations upregulate H3.3, H3.5 and H2A.J histone variants and impairs PRC2 activity by *EED* downregulation. Reduced PRC2 activity results in H3K27me3 depletion. Mutant *ATRX* (*ATRX* mut.) impairs deposition of upregulated H3.3 on telomeres. Horizontal bars in (A,F,G) indicate group mean, and the two-sided Wilcoxon test p-value is shown between groups. *MYCN*-amp, *MYCN* amplification; *TERT*-re, *TERT* rearrangement; *TERT*-high, high *TERT* expression; ALT: Alternative lengthening of telomeres. Tumors harboring more than one of the molecular characteristics are not shown in (A,B,D).

**Figure 4:**
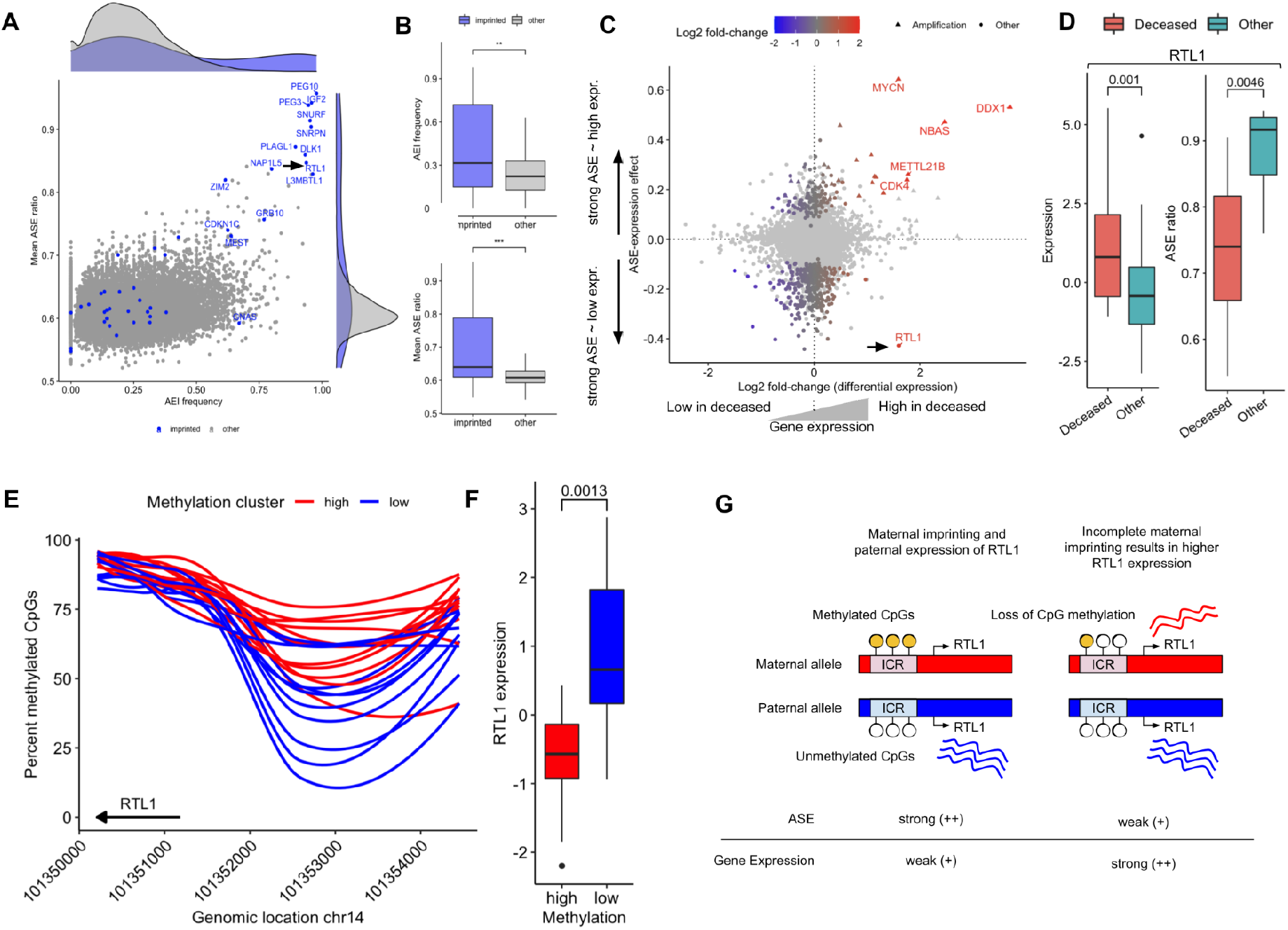
Bi-allelic expression of imprinted gene RTL1 in unfavorable tumors. **A** AEI frequency and mean ASE ratio per gene. Imprinted genes in blue, others in gray. **B** AEI frequency (top) and ASE ratio (bottom) by imprinting status per gene. **C** ASE-expression effect and differential expression (survival) per gene. Allelic regulated genes in color scale, others in light gray. **D** Gene expression (left) and ASE ratio (right) of the *RTL1* gene by survival. **E** Percent methylated CpGs in the genomic region -4 kb to +1 kb relative to *RTL1* gene start. Samples are clustered by above (high) and below (low) median methylation level in the genomic window shown. **F** *RTL1* expression by methylation clusters shown in (E). **G** Proposed model of *RTL1* upregulation through loss of maternal imprinting. Incomplete imprinting on maternal allele results in bi-allelic expression and upregulation of *RTL1*. ICR: Imprinting control region. Boxplot midlines in (B,D,F) mark median; upper and lower hinges extend to first and third quartile; upper and lower whiskers extend to the smallest and largest value max. 1.5 × IQR; p-value of two-sided Wilcoxon test is shown between groups.

Except for three tumors, *MYCN* amplifications, *TERT* rearrangements and long telomeres were mutually exclusive (Fig. 1B), in support of convergence towards a common high-risk phenotype characterized by telomere maintenance ^19–21^. MYCN amplifications were also mutually exclusive to ATRX alterations, corroborating findings on incompatibility of these two molecular traits ^38^. Comparison of *TERT* expression with telomere length estimates confirmed the existence of two distinct groups of high risk tumors: those with high *TERT* expression but short telomeres and those with low *TERT* expression but increased telomere length, indicative of ALT (Fig. 1D). In contrast, 40 of 43 low risk tumors (93%) showed neither increased telomere length (log ratio > 0.5) nor elevated *TERT* expression (z-score > -0.10, S.Fig. 5). Interestingly, active telomere maintenance was predicted in three low risk tumors (NBL09, NBL23, CB2035), which all showed disease progression. Notably, we did not find any *MYCN* amplifications in ALT samples and only a single sample with both *TERT* rearrangement and long telomeres (NBL54).

In summary, 43 tumors were classified as showing c*anonical telomere maintenance (CTM)* characterized by *TERT* activation through *MYCN* amplification, *TERT* rearrangement or high *TERT* expression. 20 tumors showed *ALT*, one tumor exhibited a *mixed phenotype (*CTM and ALT), and 51 tumors showed no signs of telomere maintenance (Fig. 1A-B, Methods). 58 of 66 high-risk tumors (88%) were classified as CTM or ALT in contrast to 7 high-risk tumors (11%) without signs of telomere maintenance. We further found a significantly reduced event-free survival for all of the three inferred mechanisms compared to tumors without telomere maintenance (*MYCN*-amp: P = 1.01 × 10^−6^; *TERT*-re: 5.56 × 10^−5^; *TERT*-high: 1.16 × 10^−7^; ALT: P = 2.47 × 10^−5^; Cox proportional hazards regression) (Fig. 1E). However, we found only *MYCN*-amp and *TERT* status, but not ALT significantly associated with overall survival (*MYCN*-amp: P = 1.17 × 10^−5^; *TERT*-re: P = 0.036; *TERT*-high: 3.16 × 10^−5^; ALT: P = 0.241; Cox proportional hazards regression) (S.Fig. 6).

### Quantifying genomic instability

We next investigated overall genomic instability in our cohort. We determined allele-specific SCNAs and overall ploidy from WGS (Methods) and classified resulting copy-number segments into states *loss, shallow loss, neutral, weak gain, medium gain, strong gain* and *focal amplification* (Fig. 2A,B) and into allelic imbalance states *balance, weak imbalance, strong imbalance, amplification* and *LOH* (S.Fig. 7, S.Methods). We additionally determined the presence of whole-genome doubling (WGD) events by phylogenetic analysis from copy-number profiles as recently described ^39^. As expected, the tumors showed pervasive patterns of genomic instability: on average 50% of the genomic regions harbored SCNAs, 31% of genomic regions showed gains and losses relative to ploidy, and 44 tumors (38%) showed WGD (Fig. 1B, S.Fig. 8). We identified gains in 17%, losses in 15% and amplifications in <0.1% of genomic regions, with distinct hotspots visible across the cohort (Fig. 2A). We found a significant enrichment of WGD in tumors without telomere maintenance (26 of 51, expected 20, P=0.03, fisher-exact test), as opposed to ALT, where fewer WGD events than expected were observed (2 of 20, expected 8, P=0.01) and tumors with canonical telomere maintenance in contrast did now show enrichment in either direction (16 of 43, expected 16, P=1.0).

Next, we determined ASE in all 115 tumors. Briefly, read counts from RNA-seq were tallied up at heterozygous germline variants (Fig. 2B) and aggregated to haplotype counts per gene using statistical phasing (S.Methods). In line with prior observations that *MYCN* amplified tumors are overall genomically more stable than their non-*MYCN* amplified counterparts ^40^ we found both the number of copy-number-imbalanced genes (P = 3.7 × 10^5^) and genes with ASE (P=0.0023) to be significantly lower in *MYCN* amplified tumors (one-sided Wilcox rank sum test) (Fig. 2C). Interestingly, we also found 4 out of 23 (17%) *MYCN* amplified tumors to harbor a substantially higher number of copy-number imbalances than the median non-*MYCN*-amplified samples (37% of genes). All 4 tumors showed signs of WGD and overall high chromosomal instability (>80%) (Fig. 2D, S.Table 1) and 3 out of 4 of these patients died from the disease. Increased genomic instability in *MYCN* amplified tumors thus might confer an additional risk factor similar to earlier findings on chromosomal losses in *MYCN* amplified tumors ^12^.

Focal amplifications were detected in 32 tumors and recurrently affected 35 genes, including COSMIC census genes ^41^ *MYCN, ALK, CDK4, LRIG3, MDM2*, and *PTPRB* (S.Fig. 9). Genes amplified in three or more tumors were exclusively detected in *MYCN* co-amplified regions on 2p24, indicating that highly recurrent amplifications are limited to the broader *MYCN* locus, likely associated with extrachromosomal circular DNAs. LOH affected 5% of genomic regions, and 1p36 LOH was found in 26 tumors (22%) of which 18 also showed amplification of *MYCN*. Shallow losses of 1p36 without LOH were detected in 6 tumors (5%). LOH of 11q was found in 42 tumors (37%), out of which 17 affected ALT tumors, 8 affected tumors with *TERT* rearrangements and 3 *MYCN*-amplified tumors. We found 11q losses without LOH in 23 tumors (20%), of which 19 were classified as shallow losses. 11q loss was found in 18 of 19 ALT tumors (90%), in line with previous reports on frequent 11q losses in ALT ^28^. Gains of 17q were highly abundant and affected 104 tumors (90%). Interestingly, 10 out of 13 strong 17q gains (78%) affected ALT tumors, suggesting that relatively higher 17q copy-numbers may be linked to the ALT phenotype in neuroblastoma. SV analysis indicated substantial heterogeneity in SV burden between tumors (S.Fig. 10A) and revealed frequent interchromosomal rearrangement between 11q13 and 17q21 (S.Fig. 10B,E) confirming earlier reports on unbalanced translocations between these chromosome arms ^42^. We analyzed the frequency of somatic SVs in 500 kb segments along the genome and detected recurrent SV breakpoints at the *MYCN, TERT, ATRX, 11q13*, and *17q21* loci (S.Fig. 10C-E), confirming previous findings on SV frequencies in a subset of tumors analyzed ^19^.

To investigate the effect of SCNAs on patient survival systematically we associated allelic copy-number imbalances on the level of chromosome arms and in non-overlapping 5Mb bins with mortality (S.Methods) and found expected associations at 1p and the *MYCN* locus as well as a yet undescribed association of 17p imbalance (S.Fig. 11A-C). Five tumors of deceased patients harbored extreme copy-number imbalances (> 0.9) due to loss of 17p (S.Fig. 12A), pointing towards elevated risk conferred through chromosomal loss. However, also 10 out of 26 donors (38%) with tumors harboring imbalanced gains died from the disease. We compared survival probabilities using the Kaplan-Meier method and found that survival was significantly reduced for tumors with 17p imbalance (P = 5.2 × 10^−4^) (S.Fig. 12B). Similarly, Cox proportional hazard regression showed that 17p imbalance is significantly associated with mortality (P = 1.44 × 10^−5^), independent of *MYCN* amplification (P = 4.32 × 10^−6^) (S.Fig. 13). Notably, 17p LOH is frequent in neuroblastoma cell lines ^43^, but its occurrence in primary neuroblastoma is less well described. Interestingly, we did not find *TP53* missense mutations or SVs, suggesting that 17p loss might act through down-regulation of neuronal genes (S.Fig. 12C-D, S.Table 2) or through a second hit in *TP53* that occurred after the sampling time point.

### Copy-number dosage is an important genetic determinant of gene expression

We next sought to identify the effects of genetic aberrations and to quantify their contribution to gene regulation in neuroblastoma. We used linear models to predict both total gene expression and the ASE ratio per gene from its lead *cis*-eQTL variant, proximal SV breakpoints, copy-number status and local mutational SNV burdens in promoter and gene body (see Methods and ^35^). For ASE analysis, an average of 5,768 (2,691–7,544) expressed genes were considered per tumor. In keeping with the literature, we found SCNAs to have the strongest effect among all genetic factors on both ASE and total gene expression ^44^, explaining an estimated 30.3% and 8.0% of variance in ASE and total expression respectively (Fig. 2E-F), and demonstrating a clear allele-specific copy-number dosage effect on gene expression on average (Fig. 2G-H). Lead germline cis-eQTL variants were the second largest genetic contributor explaining 1.6% of variance in ASE and 2.6% of variance in total gene expression. Despite emerging evidence of targeted cis-deregulation in neuroblastoma ^19,30,45^, overall somatic SVs and SNVs explain the least amount of variance in ASE and total expression with less than 1.0% and 1.2% respectively on average, in line with recent findings in adult tumors ^35^.

Even though SCNAs exhibit a strong allelic dosage effect on gene expression, transcription levels of genes are subject to transcriptional adaptations and buffering ^46,47^. To investigate dosage sensitivity in our cohort systematically, we examined copy-number components in our linear models and found statistically significant copy-number effects that explain between 2.4% to 71.0% of observed variance in gene expression (S.Fig. 14). We ranked all protein-coding genes by expression variance explained and tested for pathway enrichment using gene set enrichment analysis (GSEA, S.Methods). We found 69 Reactome pathways enriched (FDR < 0.05) for copy-number dosage effects (S.Table 3), of which 25 remained after accounting for overlapping gene sets (S.Fig. 15). Notably, dosage sensitive genes were enriched in pathways involved in cell cycle and DNA repair, and in regulation of tumor suppressor genes *TP53, PTEN and RUNX3*. In contrast, conducting the same GSEA analysis on genes ranked by total copy-number alone did not yield any significant pathway enrichment.

Copy-number dosage analysis of the *TERT* locus revealed that in CTM tumors (*MYCN*-amp, *TERT*-re, *TERT*-high) *TERT* dosage is significantly correlated with *TERT* expression (Fig. 3A, Spearman’s r = 0.63, P = 8.5 × 10^−6^) as opposed to tumors with ALT or without telomere maintenance (r = 0.15, P= 0.22), independent of sample purity (P = 0.48, ANOVA F-statistic). Our findings show that SCNAs adjust the regulatory landscape of neuroblastoma towards dysregulation of key cancer pathways and that copy-number gains effectively upregulate *TERT* in tumors with CTM (Fig. 3B), with the highest telomerase expression found in tumors with both *TERT* activation and copy-number gains.

### 11q loss and 17q polysomy link alternative lengthening of telomeres to upregulation of histone variants

To investigate if SCNAs are linked to increased telomere length in ALT tumors, we tested each chromosome arm for association between tumor DNA content and the ALT phenotype using logistic regression, controlling for *ATRX* mutations (Methods). We found 11q losses (P = 4.83 × 10^−7^, ANOVA Chi-squared test) and 17q gains (P = 2.88 × 10^−5^, ANOVA Chi-squared test) to be significantly associated with ALT (Fig. 3C), confirming previous observations of frequent 11q loss in ALT ^28^ and revealing a yet undescribed association of 17q gain with ALT. We noticed that 11q loss co-occurs with strong 17q gains in 14 tumors and observed an overall negative correlation between DNA content of both chromosome arms across the cohort (*r* = -0.45, P = 2.01 × 10^−7^, Pearson’s *correlation*) (Fig. 3D), suggesting a genomic rearrangement involving both chromosomes. Indeed, somatic SV analysis revealed 17q to 11q translocations in 19 tumors (Fig. 3E), confirming that additional copies of chromosome arm 17q translocate to 11q in the aberrant tumor karyotype ^42^. Notably, 17q gains were identified in 105 of 115 tumors (91%) independent of ALT. However, ALT tumors were significantly enriched in the strongest 17q copy-number gains (S.Fig. 16).

To pinpoint candidate genes contributing to ALT we tested for differential gene expression between ALT and non-ALT tumors, while controlling for *MYCN* amplification status, the presence of *ATRX* mutations and the sex of the patient (Methods). We found 293 such genes (FDR 0.05), of which 143 and 150 were up- and down-regulated respectively (S.Fig. 17, S.Table 4). We hypothesized that a subset of these genes might be driven by the ALT-associated SCNAs on 11q and 17q. Correlation between gene expression and DNA dosage of these chromosome arms revealed up-regulated histone variant genes *H3F3B* (17q), *H2AFJ* (12p) and *H3F3C* (12p) among genes strongly affected by 17q and 11q dosage (Fig. 3F). *H3F3B* (and its paralog *H3F3A*) encode the histone variant H3.3 ^48^, which is altered by activating mutations in several pediatric tumor entities, including tumors of the central nervous system ^49,50^ and up to 95% of chondroblastomas ^51^. Interestingly, activating H3.3 mutations triggered ALT in pediatric high-grade glioma regardless of *ATRX* mutation status ^52^, indicating that similarly, H3.3 upregulation may have functional implications in ALT neuroblastomas. *H3F3C*, which encodes for histone variant H3.5 is frequently mutated across different pediatric brain tumors, where alterations were found to be mutually exclusive to those in *TP53* and associated with reduced genome stability ^53^. The *H2AFJ* gene encodes for histone variant H2A.J and is deregulated in melanoma ^54^, breast cancer ^55^ and colorectal cancer, where its upregulation is associated with poor survival ^56^. Taken together, these results suggest that copy-number alterations may deregulate histone variants contributing to epigenetic dysregulation and genome integrity in ALT neuroblastomas.

The genetic effects model (Methods) predicted 41% and 60% of expression and ASE variance of *H3F3B* explained by local copy-number effects, indicating that expression of *H3F3B* is directly associated with 17q dosage (S.Fig. 18). However, only 3% of *H2AFJ* and 2% of *H3F3C* expression variance is explained by local copy-number effects on 12p, indicating that here ALT-associated upregulation may result from regulatory effects in *trans*.

To obtain a quantitative understanding how expression of the identified histone variant genes relates to ALT we predicted presence of ALT from the expression of *H3F3B, H3F3C* and *H2AFJ* using logistic regression. We found expression of *H3F3B* and *H2AFJ*, but not *H3F3C* to be significantly associated with ALT in the presence of the two other genes (*H3F3B*: P = 0.001; *H2AFJ*: P = 0.008 ; *H3F3C*: P = 0.543; ANOVA), suggesting that expression of *H3F3B* and *H2AFJ* is independently associated with ALT. For an independent validation, we compared the expression levels of *H3F3B* and *H2AFJ* between 130 telomeric c-circle positive and negative neuroblastomas from Hartlieb et al. ^28^, and found significantly higher expression of *H3F3B* (P = 3.01 × 10^−4^, ANOVA) and *H2AFJ* (P = 0.02, ANOVA) in c-circle positive tumors, confirming their upregulation in ALT (S.Fig. 19).

Despite *ATRX* alterations being significantly associated with longer telomeres, we did not find *ATRX* to be differentially expressed between ALT and non-ALT (S.Table 4). We speculated that interaction partners of *ATRX* could be subject to deregulation in ALT tumors. To identify potential interactions of *ATRX* and identified histone variants with proteins of differentially expressed genes in ALT we obtained direct (first order) predicted protein interactions between *ATRX, H3F3B, H2AFJ, H3F3C* and other proteins of differentially expressed genes in ALT affected by 11q or 17q dosage (S.Methods). The resulting network predicted high-confidence direct interactions between *ATRX* and differentially expressed histone 3.3 variant gene *H3F3B*, as well as *RAD51C* and *SRSF1* (Fig. 3G). A network module containing *H3F3B, H2AFJ* and *H3F3C* also included deregulated histone methylation factors *EED* and *KMT2A. EED* is part of the polycomb repressive complex 2 (*PRC2*), which modulates transcriptional repression by methylation of H3 histones ^57,58^, and we found *EED* to be down-regulated in ALT tumors by 11q-dosage effects (Fig. 3H, S.Fig. 20, S.Table 4). The PRC2 complex is frequently inactivated by *EED* loss in malignant peripheral nerve sheath tumors ^59^ and adenosquamous lung tumors ^60^. Upregulation of H3.3 and H3.5 histones and concomitant downregulation of *EED* in ALT point towards a relative depletion of H3K27me3 as a consequence of higher H3 variant histone availability and impaired PRC2 activity (Fig. 3H), similar to PRC2 inhibition by activating H3.3.pK27M mutations in pediatric gliomas ^61–63^ or expression of PRC2 inhibitor *EZHIP* in ependymomas ^64^. Investigating this in our cohort, we did neither find H3.3.pK27M nor ALT-associated upregulation of *EZHIP* (S.Fig. 21) in any of the tumors.

Our findings implicate 11q loss and strong 17q gain in ALT neuroblastomas and show that these alterations deregulate *ATRX* interaction partners. They highlight histone variants as key components of ALT-deregulated *ATRX* protein interactions and indicate that activity of the PRC2 complex could be reduced due to attenuated *EED* expression resulting from 11q loss, providing additional evidence for histone-dependent chromatin deregulation by copy-number dosage in ALT neuroblastomas.

### Imprinted RTL1 is upregulated by bi-allelic activation in unfavorable tumors

Finally, we characterized genes by ASE frequency and average ASE ratio across tumors. The highest ASE ratio (0.96) and frequency (0.98) was found for the *PEG10* gene, which is maternally imprinted in most tissues ^65^. Generally, imprinted genes ^66^ including *IGF2, DLK1, RTL1* and *L3MBTL1* (Fig. 4A) were enriched among the genes with strongest (P = 3.0 × 10^−6^) and most frequent ASE (P = 0.003, both Wilcoxon rank sum test) (Fig. 4B), showing that expression imbalance recapitulates imprinting in neuroblastoma.

Since ASE can be caused by either up- or downregulation of gene expression on one parental haplotype, we systematically explored effect directionality by testing for association between ASE and total expression. 10,862 genes that were informative for ASE in at least 20 samples were considered, out of which 455 showed a significant (FDR < 0.05) effect of ASE on total gene expression (S.Methods, S.Table 5). To narrow the search, we intersected these 455 genes with those differentially expressed between deceased and other patients, resulting in a final set of 107 candidate genes (S.Table 5). Among these, genes contained on the MYCN amplicon *MYCN, NBAS* and *DDX1* showed a positive ASE-expression effect due to strong upregulation by mono-allelic amplifications. In contrast, chromosome arm 1p (56%) and 17p (12%) were most frequent among all 76 genes with negative ASE-expression effect, indicating that loss of 1p and 17p underlies downregulation of these genes in tumors of deceased patients.

Interestingly, a substantial negative ASE-expression association was found in the Retrotransposon Gag Like 1 (*RTL1*) gene, which was upregulated in tumors of deceased patients (Fig. 4C,D). *RTL1* is a maternally imprinted gene involved in placental/neonatal development ^67^ and widely expressed in the nervous system ^68^. Upregulation of *RTL1* confers selective growth advantage in hepatocarcinoma ^69^ and promotes cell proliferation by regulating Wnt/β-Catenin signaling in melanoma ^70^. *RTL1* was one of 16 genes informative for survival time in a previous study of high-risk neuroblastomas, with stronger *RTL1* expression associated with shorter survival ^71^. Our linear model revealed only a minor contribution of SCNAs and germline variants to ASE in *RTL1* (S.Fig. 22), suggesting that differences in allelic expression levels may result from methylation differences. Analyzing a subset of tumors using Bisulfite sequencing (BS-seq) (S.Methods), we found that decreased methylation levels at CpGs upstream of *RTL1* are associated with higher *RTL1* expression (Fig. 4E,F, S.Fig. 23). Taken together these findings suggest that upregulation of *RTL1* in neuroblastoma is induced by bi-allelic activation in unfavorable tumors, likely due to loss of imprinting on the maternal allele (Fig. 4G).

## Discussion

We here systematically characterized the effects of copy-number dosage on neuroblastoma gene expression and demonstrated how copy-number gains interact with upregulated *TERT* to increase the efficacy of canonical telomere maintenance. We found 11q loss and strong 17q gain as markers of ALT in addition to *ATRX* alterations, and revealed upregulation of histone variant genes *H3F3B, H3F3C* and *H2AFJ*. Histone variants replace replication-dependent canonical histones in nucleosomes during the cell-cycle, affecting chromatin organization at telomeric ^72^ and actively transcribed regions by replication-independent chromatin incorporation ^73–75^ and interaction with chaperones and chromatin factors ^76^. *H3F3B* resides on 17q, and our findings strongly suggest that *H3F3B* is directly upregulated by 17q gains, which have already been reported to exert oncogenic effects through increased gene dosage ^43^. In contrast, *H3F3C* and *H2AFJ* expression are associated with 11q loss and 17q gain, but neither of these reside on these chromosome arms, suggesting that regulatory effects in *trans* underlie this association.

Possibly, this way copy-number alterations mediate histone replacement and chromatin re-organisation in ALT, leading to decondensation and increased transcription ^73,74^. Dosage-dependent down-regulation of the repressive PRC2/*EED*-*EZH2* complex, which methylates the lysine 27 residue of H3 histones may contribute to this reprogramming, and we found *EED*, which is predicted to interact with all three histone variants, as differentially down-regulated in ALT tumors by 11q loss. Similarly, PRC2 activity in pediatric high-grade glioma is impaired by H3.3K27M mutations altering *EZH2* binding ^63^ and resulting in depletion of H3K27 di- and tri-methylation ^62^.

*ATRX* stabilizes telomeres through depositing of H3.3 histones, thereby preventing replication-induced breaks conducive to ALT ^72,77^. In contrast, *ATRX* is not required to deposit H3.3 histones in actively transcribed regions ^72^. Consequently, H3.3 upregulation through *H3F3B* dosage in ALT tumors with defective *ATRX* may increase the prevalence of H3.3 in nucleosomes of active chromatin without its stabilizing effect at telomeres. Importantly, we found 11q loss and 17q gain to be associated with ALT independent of *ATRX* mutations. Because not all ALT tumors harbor *ATRX* alterations, deregulated histone variants may contribute to the ALT phenotype more directly. In high-grade gliomas ALT frequently occurs in H3.3G34R-mutant tumors independent of *ATRX* alterations ^52^, indicating a functional link between impaired H3.3 function and ALT.

Additionally, loss of *ATRX* alone may not be sufficient to induce ALT ^77^, and *ATRX* mutations are likely not the only molecular feature responsible for this phenotype. However, in *ATRX-*wildtype ALT-positive neuroblastomas, *ATRX* protein levels were found to be significantly decreased ^28^, suggesting that impaired *ATRX* activity could still underlie ALT in these cases. Furthermore, not all ALT-positive tumors showed 11q loss and strong 17q gain and these alterations were also present in a few ALT-negative tumors. Additional research with larger cohorts will be needed to characterize this relationship further.

We also found that 17p imbalance is associated with poor outcome in neuroblastoma. In tumors with a 17p LOH event, loss of function of *TP53* due to a second hit could be responsible for this, but no second hit was found in our cohort and we did not observe a copy-number dosage effect on *TP53* expression. Alternatively, dosage-dependent down-regulation of other genes than *TP53* on 17p could underlie this association. Survival-associated 17p copy-number dosage effects were enriched for neuronal genes, which suggests that impairment of neuronal processes could result in a more aggressive phenotype. The exact mechanism that underlies higher mortality of donors with 17p imbalance still needs to be investigated, such as the neuronal differentiation state of 17p loss or the mutational status of *TP53* in relapsed tumors that initially showed a heterozygous deletion at diagnosis.

Lastly, we identified *RTL1* as a candidate marker for unfavorable tumors due to loss of imprinting of the maternal allele, similar to earlier reports on loss of imprinting of the *IGF2* gene in Wilms’ tumors ^78^. The transposon-derived *RTL1* gene ^79^ is part of a broader imprinted *DLK1*-*DIO3* gene cluster with three paternally expressed genes *DLK1, RTL1* and *DIO3. DLK1* expression in neuroblastoma cell lines is associated with neuroendocrine lineage differentiation ^80^. Possibly, imprinting heterogeneity at the *DLK1*-*DIO3* gene cluster reflects differentiation states in neuroblastoma progenitor cells, and incomplete imprinting may characterize the cell-of-origin of more aggressive or treatment-resistant tumors.

Our analyses shed light on the complex interaction of genetic, epigenetic and transcriptomic effects and how gene dosage interacts with other genetic and epigenetic factors to shape the regulatory landscape of neuroblastoma. In the future, the increase in size of such multi-omics datasets will enable a more complete understanding of the development of disease-relevant phenotypes and potentially convergent pathways.

## Methods

### Whole-genome- and RNA-sequencing data

This study makes use of sequencing data from tumor and blood samples of patients diagnosed with neuroblastoma that were enrolled and treated according to trial protocols of the German Society of Pediatric Oncology and Hematology (GPOH) in the multi-center study GPOH-NB2004 ^34^. This study was conducted in accordance with the Declaration of Helsinki and Good Clinical Practice, and informed consent was obtained from all patients or their guardians. Collection and use of patient specimens was approved by the institutional review boards of Charité Universitätsmedizin Berlin and of the Medical Faculty, University of Cologne. Collected specimens and clinical annotations were made available by Charité-Universitätsmedizin Berlin or the National Neuroblastoma Biobank and Neuroblastoma Trial Registry (University Children’s Hospital Cologne) of the GPOH. Corresponding sequencing data is available from the European Genome-phenome Archive (https://www.ebi.ac.uk/ega/) under accession number EGAS00001001308. Analyzed DNA and RNA samples were obtained from primary tumors of at least 60% tumor cell content as evaluated by a pathologist. *MYCN* copy-number was determined by FISH. Sample preparation and sequencing of DNA and RNA was performed as described earlier ^8^. WGS of tumor-normal pairs was performed on the HiSeq X-Ten platform (Illumina, San Diego, USA), yielding paired-end reads of 2×150 bp length. Ribo-depleted RNA was sequenced on the HiSeq4000 platform (Illumina, San Diego, USA) yielding reads of 2×150 bp length. Additional sequencing data was obtained from the European Genome-phenome Archive under accession number EGAS00001001308 for a non-overlapping set of donors from a previous study on somatic structural rearrangements in neuroblastoma ^19^. After quality control 52 donors of this study were included, yielding a total of 115 donors with matched tumor RNA-seq, tumor WGS, and blood-derived normal WGS.

### Telomere maintenance analysis

Telomere lengths were estimated from WGS of normal and tumor samples by Telseq 0.0.2 ^37^ with parameter -u (ignore read groups) and otherwise default settings. Briefly, the method estimates telomere length by counting WGS reads containing the telomere repeat sequence (TTAGGG)^*k*^, where *k* denotes the number of repeats of the 6-mer. Telseq uses default repeat length k = 7 and normalizes the resulting read count by GC content and a genome size factor. The authors calibrated the default parameters using telomere length measurements determined by southern blot analysis of terminal restriction fragments. We summarized telomere lengths per sample by the log telomere length ratio log(TLR) = log(L_T_/L_N_), where L_T_ and L_N_ are the Telseq estimates for telomere length in tumor and normal WGS sample respectively.

Tumors were clustered based on unsupervised gaussian mixture modeling of *TERT* gene expression (N=2 mixture components). A threshold (z-score > -0.1028) for high *TERT* expression was defined as the *TERT* expression z-score at which the probability of assignment of a tumor to the component of stronger *TERT* expression exceeded 95%, similarly as described in ^20^. We assigned the *TERT-high* attribute to tumors that exceeded the *TERT* expression threshold and for which neither *MYCN* amplification nor *TERT* rearrangements were detected.

The telomere maintenance status (CTM, ALT, Mix, or None) was assigned based on the status of *MYCN* amplification, *TERT* rearrangement, *TERT-high* attribute and telomere length ratio as follows: Status *canonical telomere maintenance* (CTM) was assigned to tumors with either *MYCN* amplification, *TERT* rearrangement or that were classified as *TERT-high*. Status *alternative lengthening of telomeres* (ALT), was assigned to tumors with telomere length ratio log(L_T_/L_N_) > 0.5. Status *Mix* was assigned to all tumors that met criteria for both CTM and ALT. Status *None* was assigned to all other tumors.

### Allele-specific copy-number analysis

Pileups of primary tumor WGS were generated by Bcftools 1.8 mpileup at SNP positions of the genotype panel established (S.Methods). Unmapped reads, or reads that were marked as optical duplicates or as “not primary alignment” were not considered in the pileup. For each of the SNPs the allelic depths were calculated from the pileups on normal and tumor alignments respectively. For SNPs with total depth of 10 or more reads in both tumor and normal alignments we determined the B-allele frequency (BAF) and the coverage log ratio (LogR). For a given pileup position the BAF is defined as the ratio between alternative allele nucleotide count and the number of total considered counts a_i_/(r_i_+a_i_), where a_i_ and r_i_ are the allelic depths of alternative and reference allele respectively. The LogR at SNP position *i* was defined as log2((d_ti_/d_ni_)/(*∂*_t_/*∂*_n_)), where d_ti_ is the total depth at SNP position *i* in the tumor sample, d_ni_ is the total depth at SNP position *i* in normal sample and *∂*_t_ and *∂*_n_ are mean depths at SNPs of tumor and normal sample respectively.

The BAF of a heterozygous SNP position is informative for the proportion of aligned reads originating from the paternal and maternal allele. At a homozygous SNP the BAF is expected to be close or equal to 1, if the sample’s SNP genotype is homozygous alternative or close or equal to 0 if the genotype is homozygous reference. The BAF is calculated separately for alignments of normal and tumor, resulting in a *normal BAF* and a *tumor BAF* per SNP and sample. The LogR is a measure of total coverage difference between normal and tumor samples and is informative at any position, including homozygous and heterozygous SNPs. It is calculated for a pair of alignments (tumor and normal), resulting in a LogR value per SNP and sample.

Allele-specific copy-number profiles were generated from tumor and normal BAFs and LogR values for each sample using ASCAT 2.6^81^ with a custom segmentation procedure. In ASCAT’s segmentation step the BAF and LogR values are converted into intervals of similar values. ASCAT’s original implementation of this segmentation considers both LogR and BAFs to obtain start and end points for segments. We found noisy coverage log ratios to introduce over-segmentation in some samples and therefore replaced the segmentation procedure with a custom implementation that only considers BAFs to determine start and end points of segments, but still estimates the segment’s coverage using the log coverage ratios. ASCAT’s output comprises copy-number segments with integer copy-numbers of major and minor alleles as well as estimates for tumor purity and ploidy. All copy-number segments were inspected manually for quality. For samples with estimated tumor purity less than 60% copy-number calling was rerun with adjusted purity and ploidy values that were manually selected after inspection of the goodness-of-fit plots and in agreement with pathology estimates of tumor purity.

Tumor purity is defined as the fraction of tumor cells in the sample or biopsy, for which the DNA was extracted. E.g. infiltration of immune cells, vascularization or capturing non-cancerous neighboring tissue in a biopsy decreases the purity of the tumor sample. Tumor ploidy describes the DNA content of tumor cells by their average haplotype count. Most healthy human cells contain two sets of chromsomes 1-22 and a pair of sex chromsomes and thefore have a ploidy of 2. In tumor cells this value can deviate due to chromosomal aberrations. Allelic gains may increase the overall DNA content and can result in ploidy values above 2. In contrast, if losses are dominating, the ploidy may be lower than 2. Because chromosomal aberrations often only affect a subset of the genome (such as individual chromosome arms or smaller regions) tumor ploidy may represent a fraction or floating point value and not necessarily an integer.

We associated the copy-number per chromosome arm with telomere length. For this purpose we derived a copy-number LogR per chromosome arm and tumor by an overlap length-weighted mean considering all copy-number segments of a given sample overlapping a chromosome arm. Samples were divided into two groups (ALT, non-ALT), excluding a single tumor (NBL54) with signs of both ALT and canonical telomere maintenance by *TERT* rearrangement (group *Mix)*. We then used this binary (ALT, non-ALT) outcome as the response of generalized linear model controlling for covariates *MYCN* amplification, *ATRX* alteration, age, sex, cohort, tumor purity, and tumor ploidy. The association p-value was determined by an analysis of variance (ANOVA) using a Chi-Squared test between two generalized linear models, of which the first modeled ALT by the covariates above and the second additionally included a term for the copy-number LogR. P-values were determined for each chromosome arm and corrected by the Bonferroni method. Chromosome arms below 0.05 FWER were considered significant.

### Gene expression analysis

Aligned tumor RNA-seq reads were counted using HTseq/htseq-count 0.9.1 on exons of protein coding genes according to Ensembl release 75 human gene annotations for the GRCh37 reference, summarizing counts on gene-level. We normalized gene expression for the purpose of between-sample comparisons in a given gene. To mitigate the effect of sequencing depths and batch effect introduced by different RNA library preparation- and sequencing methods between the two cohorts we normalized htseqs by the following strategy: We first calculated library-size normalized DESeq2 variance stabilized counts from htseq counts. Then, we modeled the variance stabilized counts by cohort membership using a linear model for each gene and determined the residual for each gene and sample. If not indicated otherwise, this residual was used as the measure for gene expression in our analyses. In addition we measured allele-specific expression (ASE) for genes with at least one expressed heterozygous SNP and sufficient coverage in a given sample (S.Methods).

We analyzed gene expression differences between ALT and non-ALT tumors by linear regression similar to the analysis that identified copy-number differences between these groups described above. We expect that this approach facilitates detection of expression differences mediated by the ALT-associated SCNAs identified. Expression values were modeled by linear combination of ALT status, *MYCN* amplification, *ATRX* alteration, age, sex, cohort, tumor purity and tumor ploidy. The p-value was derived from an ANOVA Chi-squared test for significance of the ALT status covariate and adjusted for multiple testing using the Benjamini Hochberg method. Genes with FDR < 0.05 were considered as significantly different expressed in ALT tumors.

### Analysis of genetic effects on gene expression and ASE

We modeled both total expression and ASE by local genetic effects based on detected germline and somatic variation at the respective gene locus and additional covariates using linear regression. ASE was modeled by the heterozygosity status of the SNP with greatest effect size from eQTL mapping, the copy-number ratio and binary variables indicating the presence of a structural variation breakpoint overlapping with gene coordinates including +/-100kb flanking regions, somatic SNVs in the promoter, and at gene coordinates (including UTRs and introns) as determined by Ensembl variant effect predictor (VEP) version 101.0. Similarly, gene expression was modeled by the genotype (encoded as number of alternative alleles) of the SNP with greatest effect size from eQTL mapping, copy-number LogR, somatic structural variation and somatic SNVs in promoter and gene. Tumor purity and *MYCN* amplification status were used as additional covariates in models of both expression phenotypes. In the ASE model, the log sum of coverage at the ASE SNPs was used as an additional covariate. A linear model with up to 115 observations was fitted for each gene separately. Only genes with 20 or more complete observations (for effects/covariates and expression phenotype) were considered. The explained variance per genetic effect was determined by its relative contribution to the total sum of squares as given by ANOVA on the fitted model. Significant variance components were determined by ANOVA’s F-statistic and the resulting p-value was adjusted for multiple testing by the Bonferroni method for each effect. Significant effects per gene were defined as effects at 0.05 FDR.

## Data Availability

The data analyzed in this study is available from the European Genome-phenome Archive (https://www.ebi.ac.uk/ega/) under accession number EGAS00001001308 and EGAS00001004022.

## Author contributions

M.B., A.E., J.S., S.M.W., D.B., R.F.S. and U.O. contributed to the study design and the collection and interpretation of the data. E.B. and N.T. performed quality control of sequencing data and aligned WGS and RNA-seq reads. M.B. analyzed WGS and RNA-seq data. M.B. performed allele-specific expression and allele-specific copy-number analysis. E.B., N.T and M.B. analyzed somatic single nucleotide variants. J.T. and M.B. analyzed somatic structural variants. R.M. and M.B. performed QTL mapping. M.B. conducted telomere length analysis. M.B. and C.W. conducted copy-number-survival analysis. M.B., R.F.S. and U.O. wrote the manuscript. J.S., R.F.S. and U.O. led the study design.

## Acknowledgements

This study was supported by funding from the Berlin Institute of Health project “TERMINATE-NB” to AE and UO, as well as DFG research training group “GRK 1772: Computational Systems Biology” (project number 191312833) to MB and UO. The authors would like to thank the Helmholtz Association, Germany, for support. RFS is a Professor at the Cancer Research Center Cologne Essen (CCCE) funded by the Ministry of Culture and Science of the State of North Rhine-Westphalia. The authors would like to thank Martin Peifer for helpful comments and suggestions.

