## Supplementary Material for "Copy-number dosage regulates telomere maintenance and disease-associated pathways in neuroblastoma"

### Supplementary figures

Supplementary figure 1: Risk stratification scheme of the NB2004 neuroblastoma trial

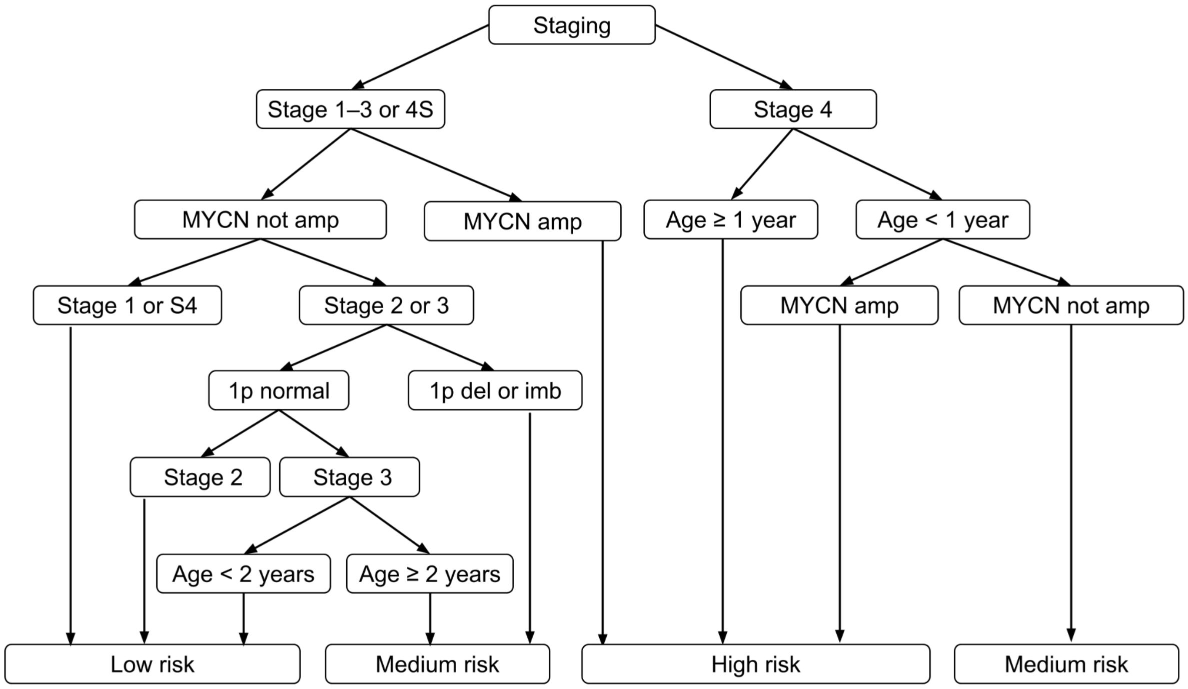

Supplementary figure 2: Frequently mutated genes in 115 primary neuroblastoma tumors by inferred telomerase maintenance mechanism

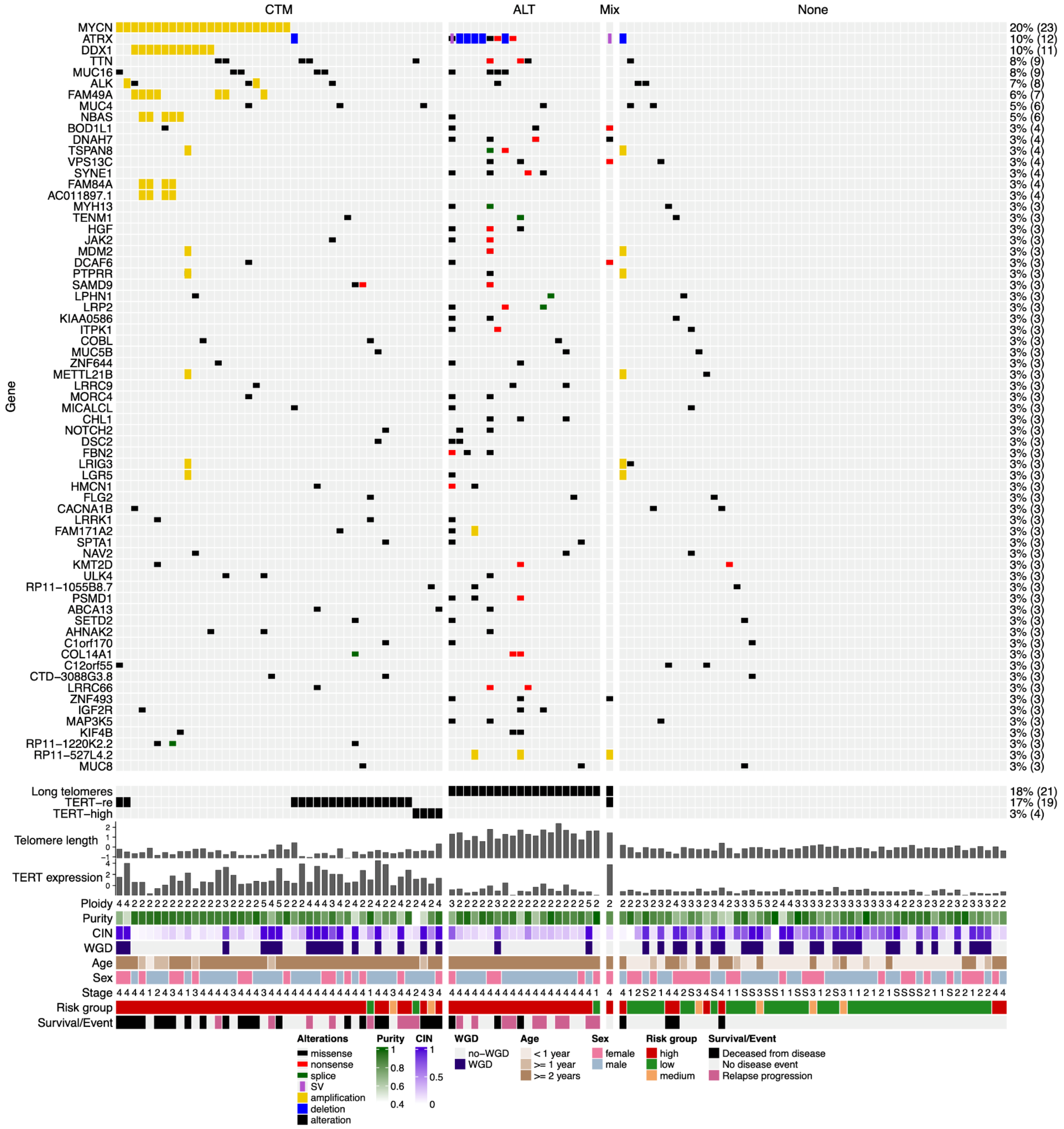


CTM, canonical telomere maintenance; ALT, alternative lengthening of telomeres; *MYCN*-amp, *MYCN* amplification; *TERT*-re, *TERT* rearrangement; *ATRX*-mut, *ATRX* mutation

Supplementary figure 3: Telomere length ratio, ALT classification and APB status

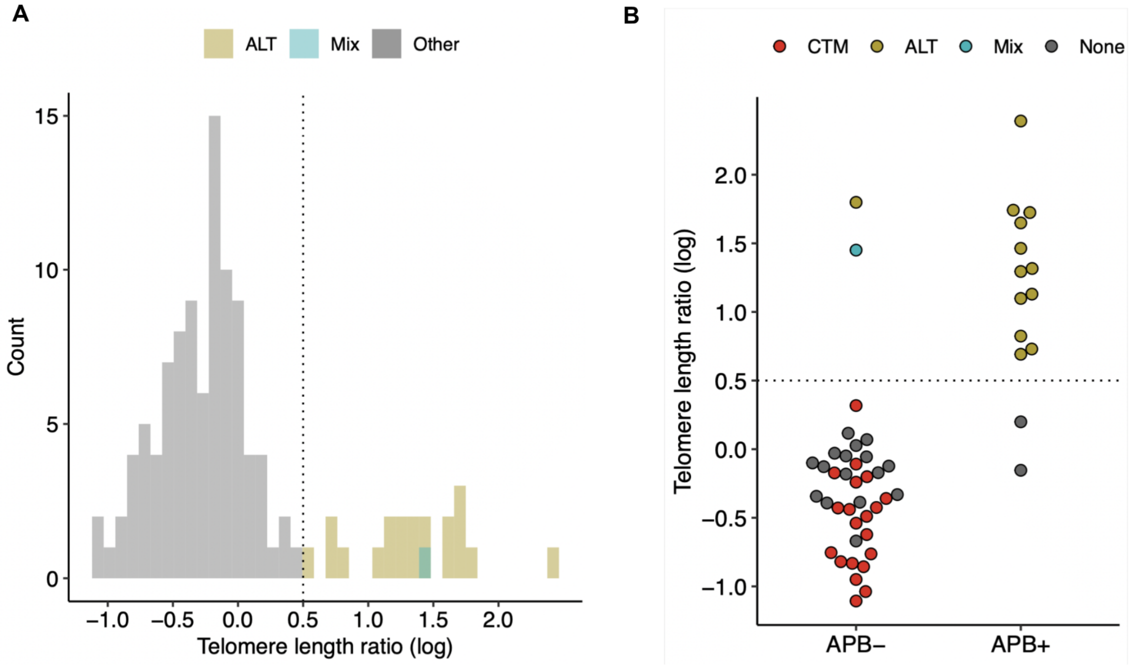


**A**, Distribution of telomere length ratio (TLR) from WGS of 115 tumors/normal pairs. Color indicates inferred telomere maintenance status “ALT” and “Mix”, other status in gray. **B**, Telomere length ratio by ALT-associated PML-nuclear bodies (APB)-status from [1] and assigned telomere maintenance group in a subset of 52 samples analyzed. Threshold [log(TLR)>0.5] as prerequisite for ALT classification shown as dotted vertical and horizontal lines in (A) and (B) respectively. A single sample (NBL54) was assigned telomere maintenance status “Mix” as it showed signs of both alternative lengthening of telomeres (ALT) and canonical telomere maintenance (CTM) by increased telomere length, *TERT* rearrangement and strong *TERT* expression.

Supplementary figure 4: Telomere length and ATRX mutation status

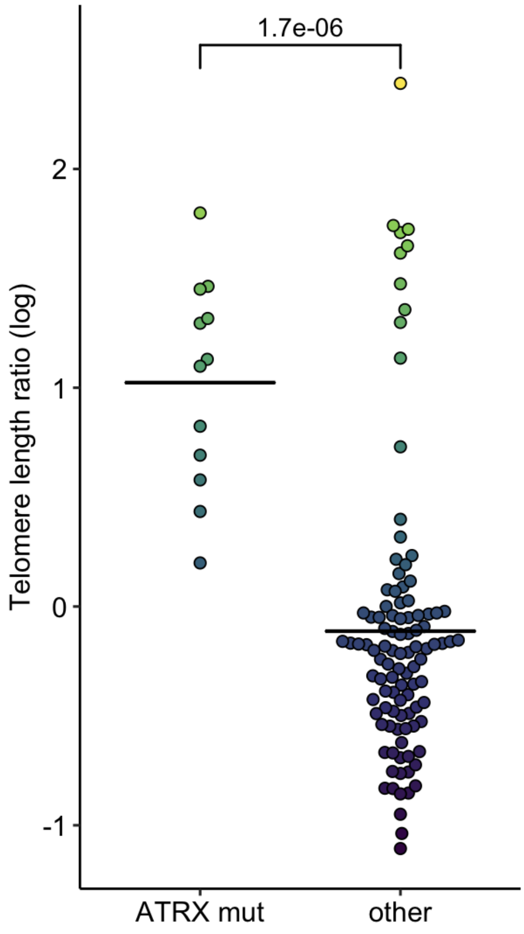


P-value between groups determined by one sided Wilcox rank sum test shown above bracket. *ATRX* mut, *ATRX* mutation.

Supplementary figure 5: Distribution of TERT expression and threshold to define high TERT expression

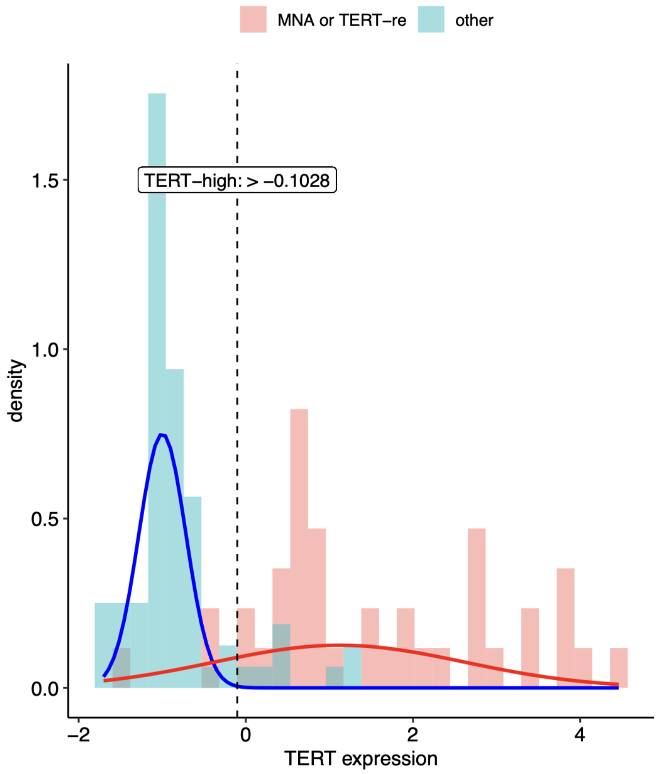


Density curves of two components derived from unsupervised gaussian mixture modeling in blue and red. Threshold for high *TERT* expression indicated as vertical dashed line. The threshold is defined as >95% probability for an observation belonging to the high expression component (red) similarly as described in ref [1].

Supplementary figure 6: Kaplan-Meier estimate of overall survival (OS) by molecular telomere maintenance features

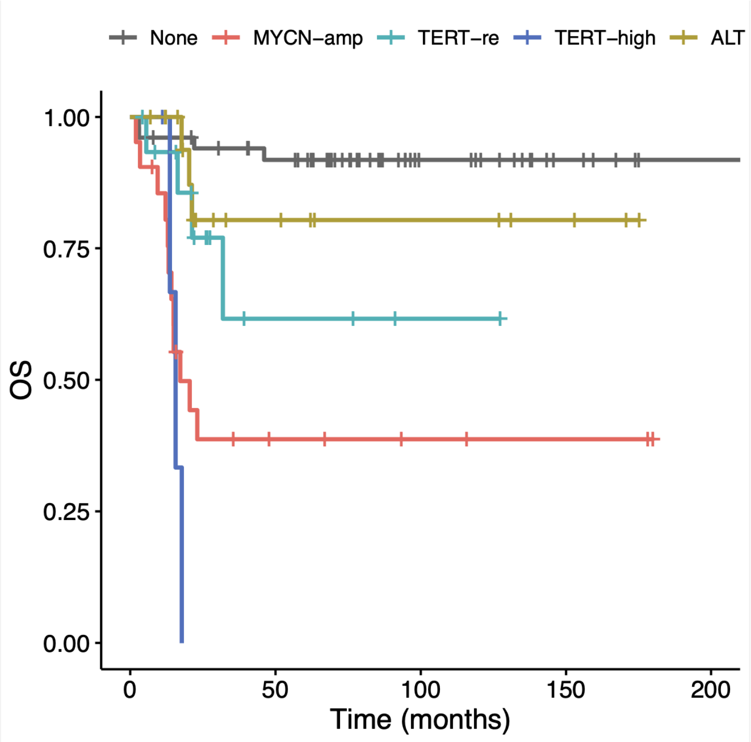


Molecular telomere maintenance features: *MYCN* amplification (*MYCN*-amp), *TERT* rearrangement (*TERT*-re), high *TERT* expression and lack of *MYCN*-amp and *TERT*-re (*TERT*-high) and alternative lengthening of telomeres (ALT). OS, Overall survival.

Supplementary figure 7: Genomic and expression imbalance

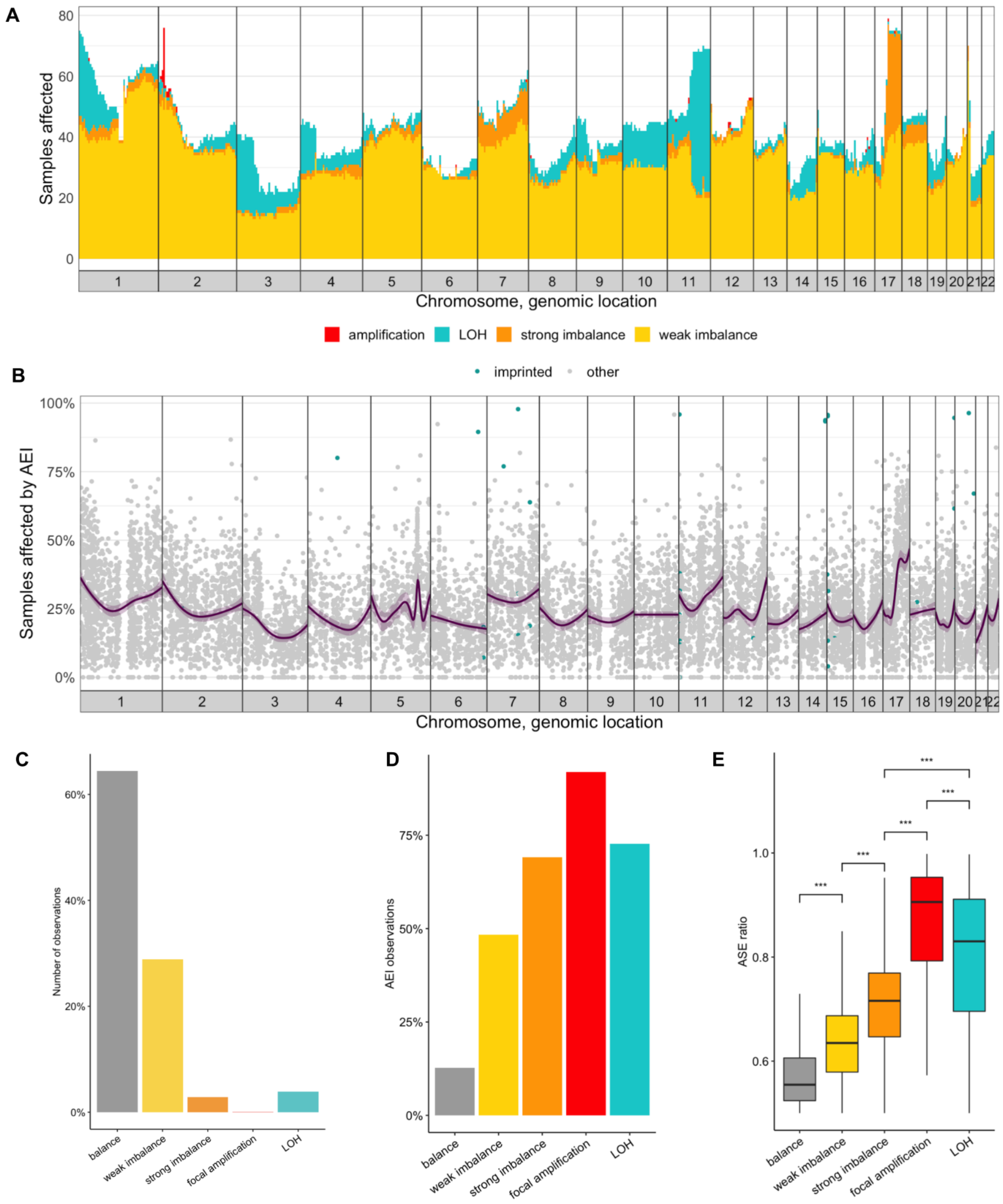


Genome-wide frequencies of somatic copy-number and allelic expression imbalance imbalances. **A**, Number of samples affected by copy-number imbalances summarized in 5Mb genomic bins. **B**, AEI frequency per gene. Dark purple line: Smoothed average AEI percentage. Light purple ribbon: 95% confidence interval of average AEI percentage. **C**, Number of observations (gene-sample pair) per copy-number balance state. **D**, Percent of observations with allelic expression imbalance. **E**, Distribution of ASE ratios, outliers not shown. Midline in boxplots marks median. Upper and lower hinges mark the first and third quartile. Upper and lower whiskers extend to the smallest and largest value max. 1.5 × IQR. AEI: allelic expression imbalance, LOH: loss of heterozygosity. ***: two-sided Wilcoxon rank sum test P < 0.001.

Supplementary figure 8: Ploidy and fraction of LOH per tumor indicating WGD status

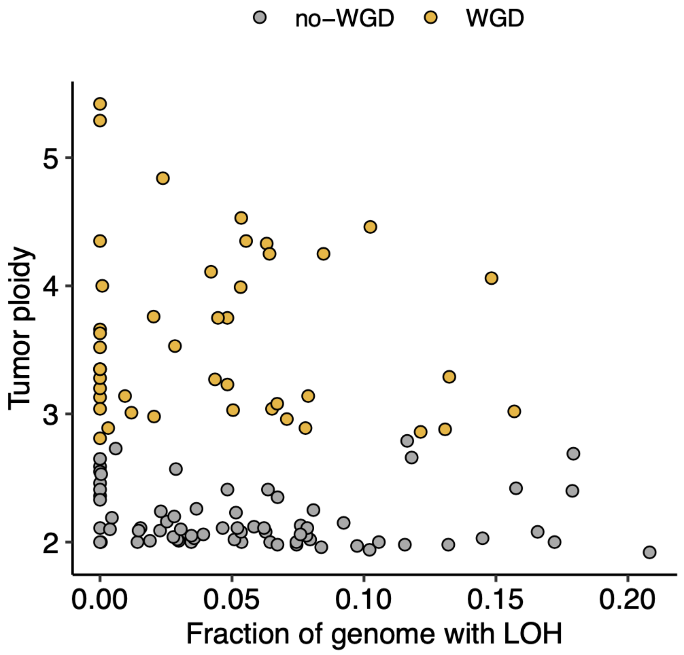


WGD, Whole-genome doubling; LOH, Loss of heterozygosity.

#### Supplementary figure 9: Recurrently amplified protein-coding genes

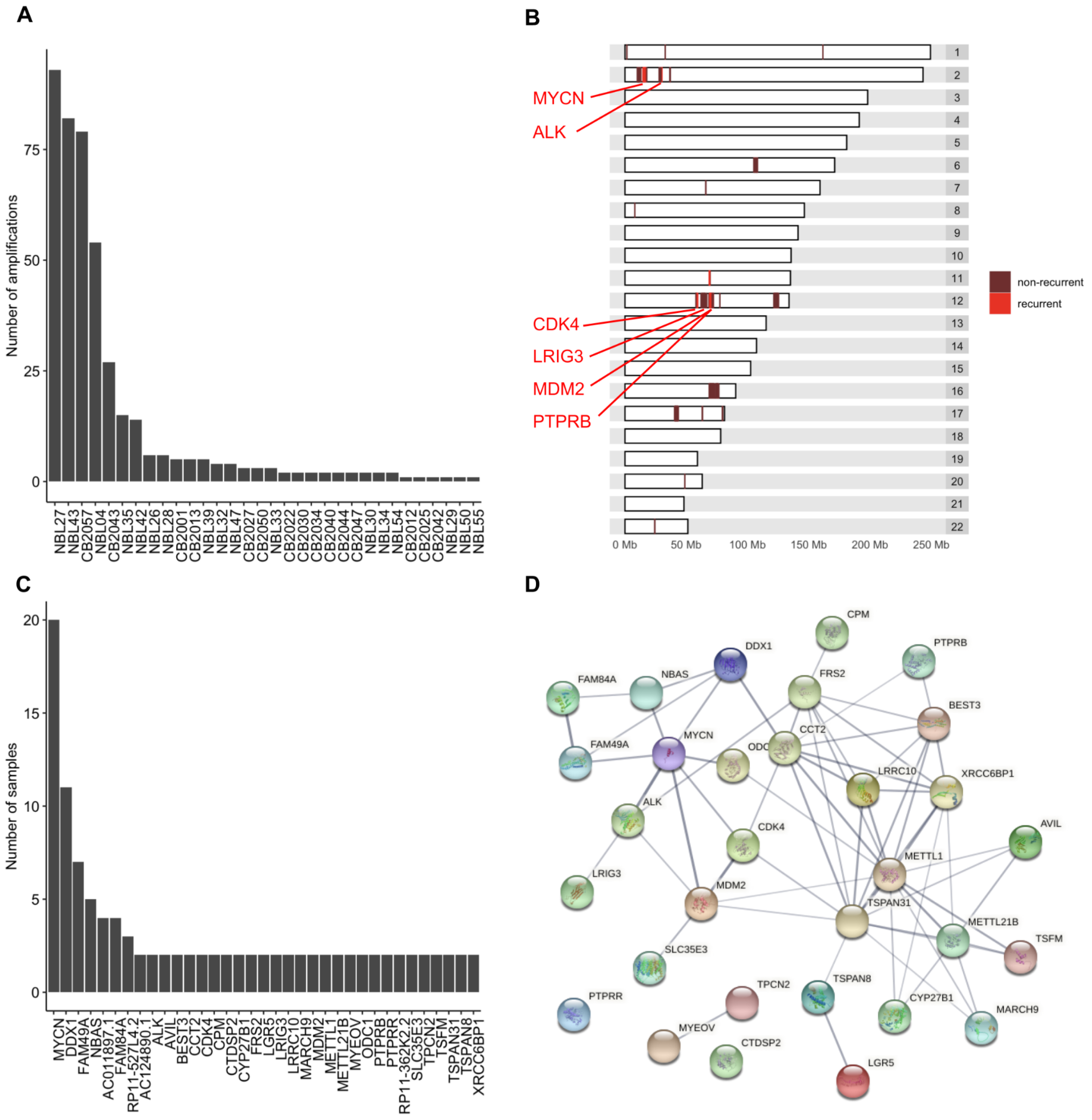


**A**, Number of amplifications per sample (samples without gene amplification are not shown). **B**, Chromosomal locations of amplified genes. Recurrently amplified COSMIC census genes are labeled by gene name. **C**, Number of samples affected by amplification in recurrently amplified genes (i.e. genes amplified in at least two samples). **D**, Network of protein interactions from the STRING database[1] for recurrently amplified genes. Thickness of edges represent data support for interactions. Network interaction enrichment $P < 1.0 \times 10^{-16}$ as given by STRING network statistics.

---

[1] https://string-db.org/, network visualization (STRING version 11.0b)

Supplementary figure 10: Genome-wide patterns of structural variation

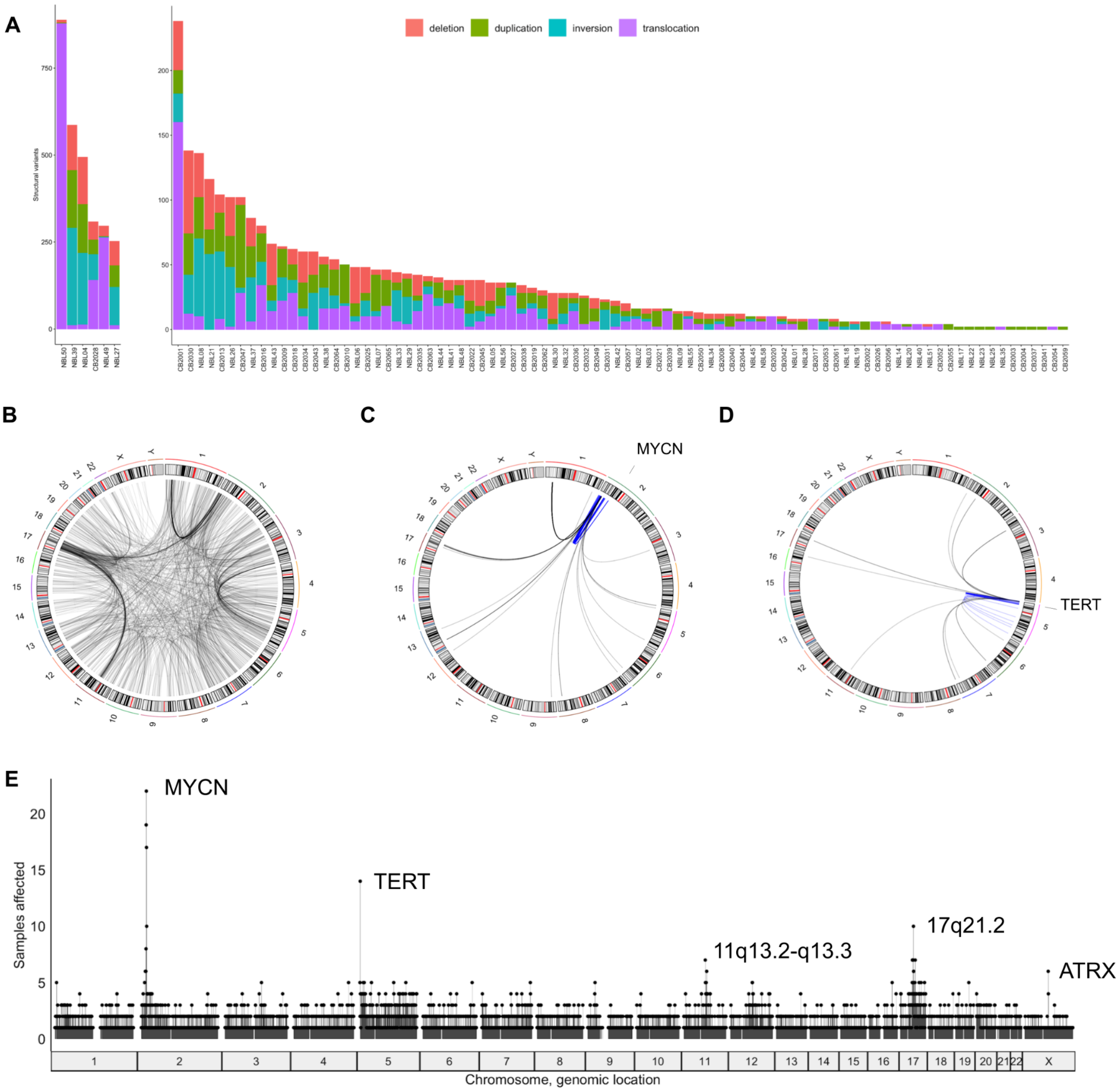


**A**, Number of structural variants per class (deletion, duplication, inversion, translocation) and sample. Samples without detected SVs not shown. **B,** Interchromosomal translocations. Structural variation with breakpoints in a 5 Mb window around *MYCN* (**C**) and *TERT* (**D**) respectively. Intrachromosomal SVs in (C,D) in blue, others in gray. **E**, Number of tumors affected by structural variation in 500 kb genomic bins.

Supplementary figure 11: Associations of copy-number ratio and disease-specific survival

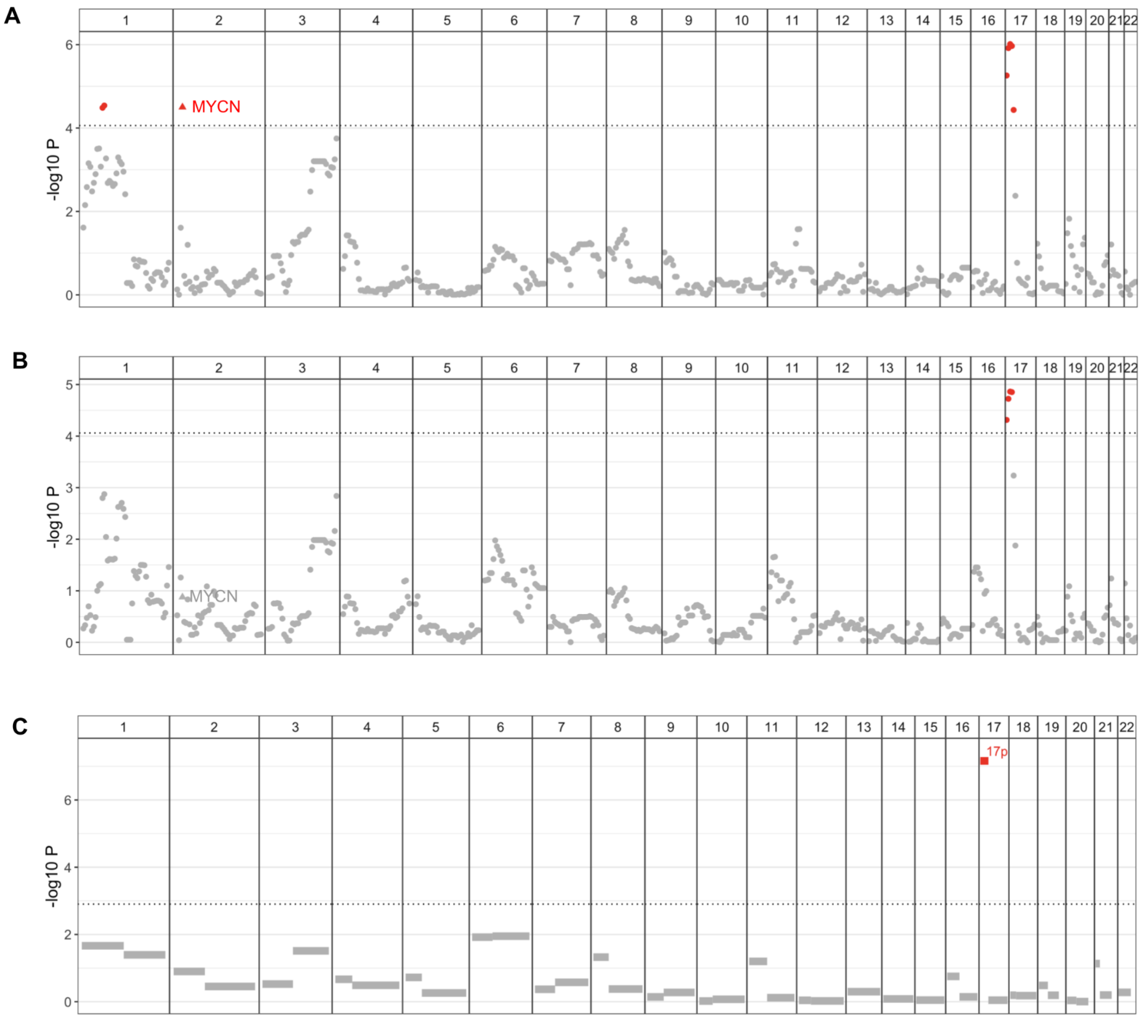


**A**, Association in 5 Mb genomic bins. **B**, Association in 5 Mb bins controlling for *MYCN* amplification status. **C** Association on the level of chromosome arms. Triangles indicate overlap between the 5 Mb genomic bin and *MYCN*. Copy significant after adjusting p-value for multiple testing in red, others in gray. Significance threshold (FWER 0.05) demarcated by horizontal dotted line.

Supplementary figure 12: 17p imbalance and copy-number dosage-dependent downregulation of neuronal genes in a subset of unfavorable tumors

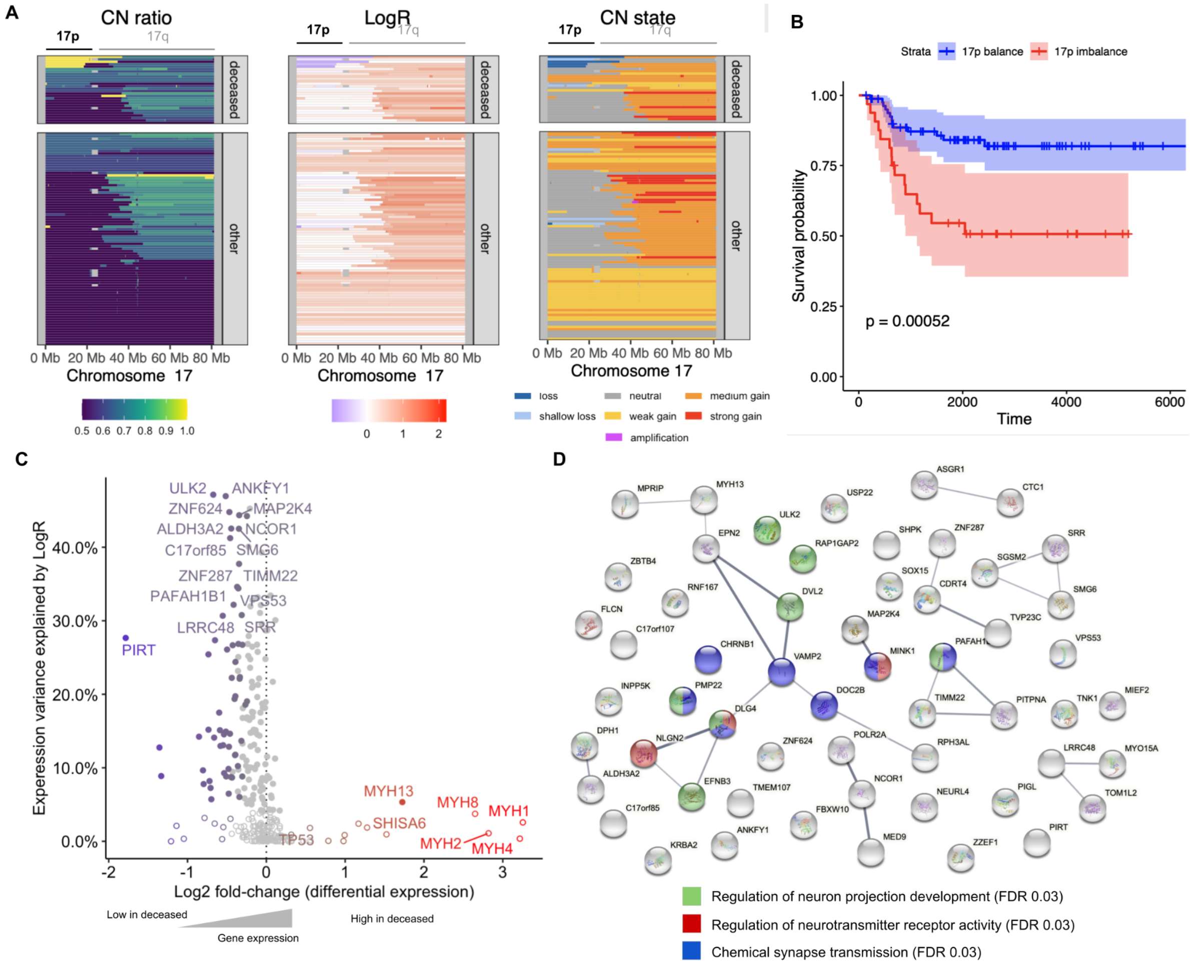


**A**, Copy-number ratio, LogR and copy-number state of copy-number segments on chromosome 17 (p and q). **B**, Kaplan-Meier estimate for survival curve by 17p copy-number imbalance. Censored data indicated by vertical marks. Colored ribbons correspond to 95% confidence intervals. **C,** Expression variance explained by LogR and differential expression Log2 fold-change. Differentially expressed genes in color scale, others in light gray. copy-number dosage effect genes as filled circles, others as empty circles. Genes of Log2 fold-change >1.5 or those >30% expression variance explained by LogR labeled by name. *TP53* is additionally labeled. **D**, Protein interaction network of differentially expressed copy-number dosage effect genes on 17p colored by selected biological processes indicated below.

Supplementary figure 13: Cox proportional hazards regression model including chromosome 17p imbalance status

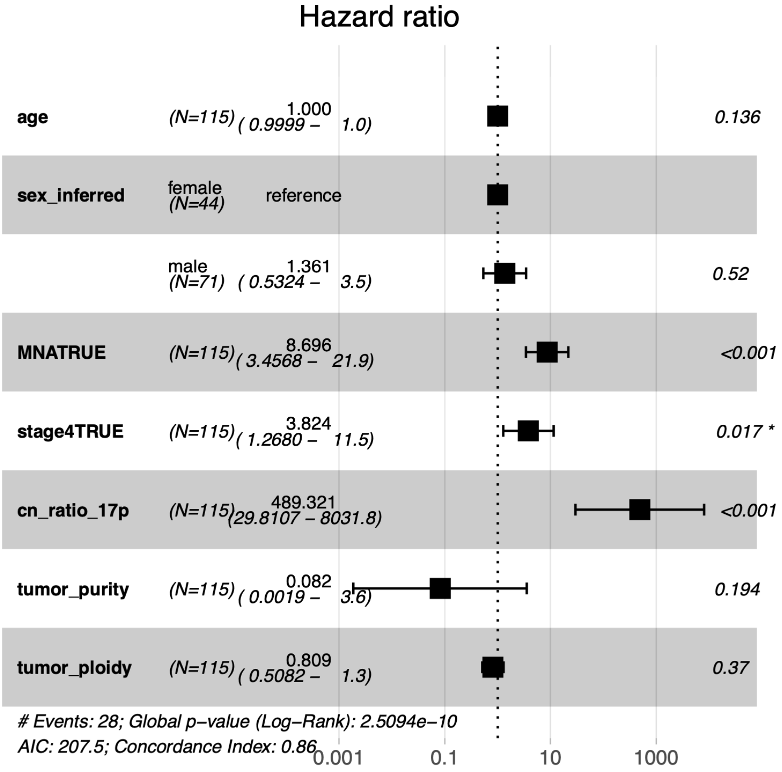

#### Supplementary figure 14: Genome-wide copy-number dosage effects

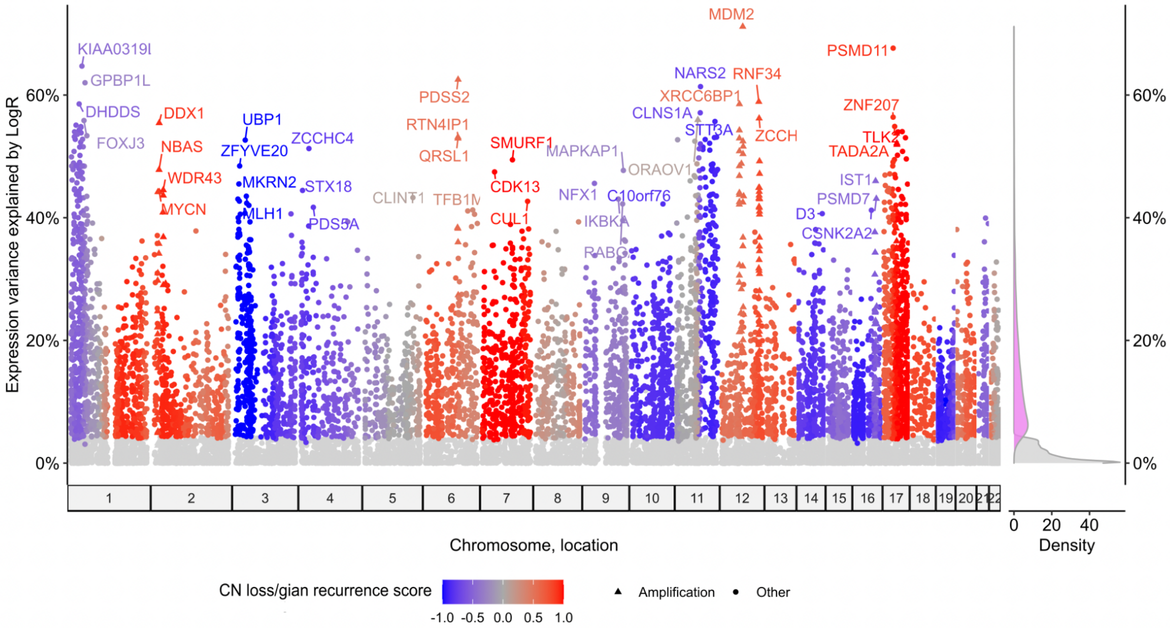


Dosage effect as variance of total expression explained by LogR per gene. Dark gray and shades of red and blue indicate significant dosage copy-number effect genes (FDR < 0.05, Benjamini-Hochberg). Non-significant genes in light gray. Color scale indicates copy-number recurrency score with shades of red and blue depicting recurrent copy-number gains and losses respectively. Genes amplified in one or more tumors as triangles. Top four genes with at least 40% variance explained per chromosome are labeled. Right: Densities of percent variance explained for significant genes (magenta) and non-significant genes (light gray).

#### Supplementary figure 15: Pathways enriched in copy-number dosage effects on gene expression

| Pathway | NES | pval | padj |
|---|---|---|---|
| Antigen processing: Ubiquitination & Proteasome degradation | 1.22 | 1.0e−05 | 8.9e−04 |
| Cell Cycle | 1.22 | 1.0e−05 | 8.9e−04 |
| Cellular responses to stress | 1.28 | 1.0e−05 | 8.9e−04 |
| Chromatin modifying enzymes | 1.29 | 1.0e−05 | 8.9e−04 |
| Deubiquitination | 1.28 | 1.0e−05 | 8.9e−04 |
| Diseases of signal transduction | 1.22 | 1.0e−05 | 8.9e−04 |
| DNA Repair | 1.24 | 1.0e−05 | 8.9e−04 |
| Gene expression (Transcription) | 1.19 | 1.0e−05 | 8.9e−04 |
| HIV Infection | 1.29 | 1.0e−05 | 8.9e−04 |
| M Phase | 1.23 | 1.0e−05 | 8.9e−04 |
| Membrane Trafficking | 1.16 | 1.0e−05 | 8.9e−04 |
| Metabolism of proteins | 1.17 | 1.0e−05 | 8.9e−04 |
| Metabolism of RNA | 1.25 | 1.0e−05 | 8.9e−04 |
| Processing of Capped Intron−Containing Pre−mRNA | 1.25 | 1.0e−05 | 8.9e−04 |
| Regulation of HSF1−mediated heat shock response | 1.45 | 1.0e−05 | 8.9e−04 |
| Transcriptional regulation by RUNX1 | 1.25 | 2.0e−05 | 1.4e−03 |
| tRNA processing in the nucleus | 1.44 | 2.0e−05 | 1.4e−03 |
| Transport of Mature mRNA derived from an Intron−Containing Transcript | 1.39 | 3.0e−05 | 1.7e−03 |
| Regulation of TP53 Activity | 1.29 | 4.0e−05 | 2.1e−03 |
| Translation | 1.20 | 5.0e−05 | 2.5e−03 |
| PTEN Regulation | 1.30 | 6.0e−05 | 2.8e−03 |
| Epigenetic regulation of gene expression | 1.30 | 9.0e−05 | 4.2e−03 |
| MAPK6/MAPK4 signaling | 1.33 | 1.0e−04 | 4.5e−03 |
| Transport of Ribonucleoproteins into the Host Nucleus | 1.47 | 1.6e−04 | 6.5e−03 |
| Transcriptional regulation of white adipocyte differentiation | 1.32 | 2.0e−04 | 7.4e−03 |

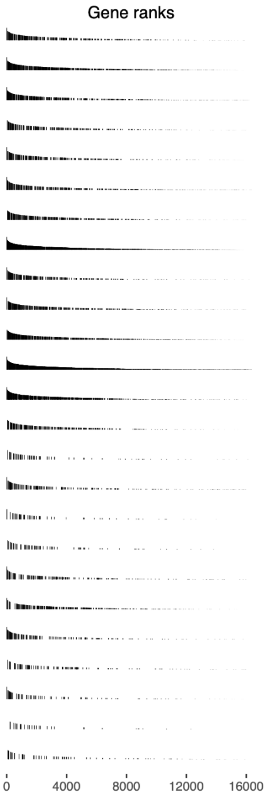


Independent reactome pathways enriched for copy-number dosage effect on gene expression. Black bars in column “Gene ranks” indicate pathway membership of genes ranked by copy-number dosage effect. NES, normalized enrichment score.

Supplementary figure 16: Log-ratio of 17q coverage difference between tumor and normal sample by alternative lengthening of telomeres status

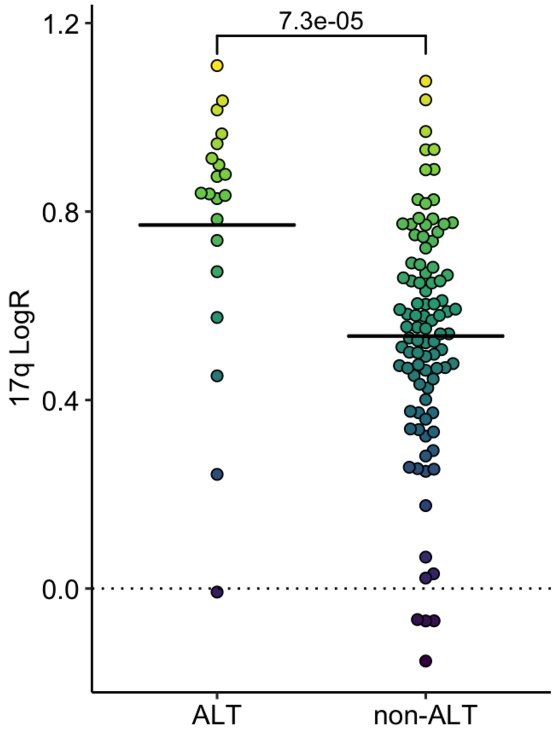


P-value between groups determined by one sided Wilcox rank sum test. ALT, alternative lengthening of telomeres.

Supplementary figure 17: Differential expression analysis between ALT and non-ALT tumors

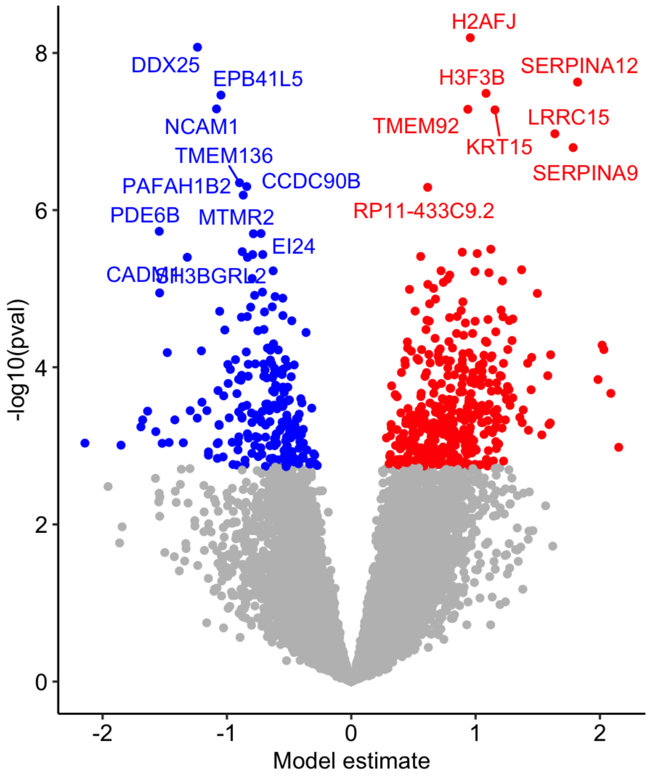


Red indicates upregulated genes in ALT, blue indicates ALT down-regulated genes (FDR < 0.05). ALT, alternative lengthening of telomeres.

Supplementary figure 18: Local genetic effects on expression and ASE in H3F3B, H2AFJ, H3F3C and EED

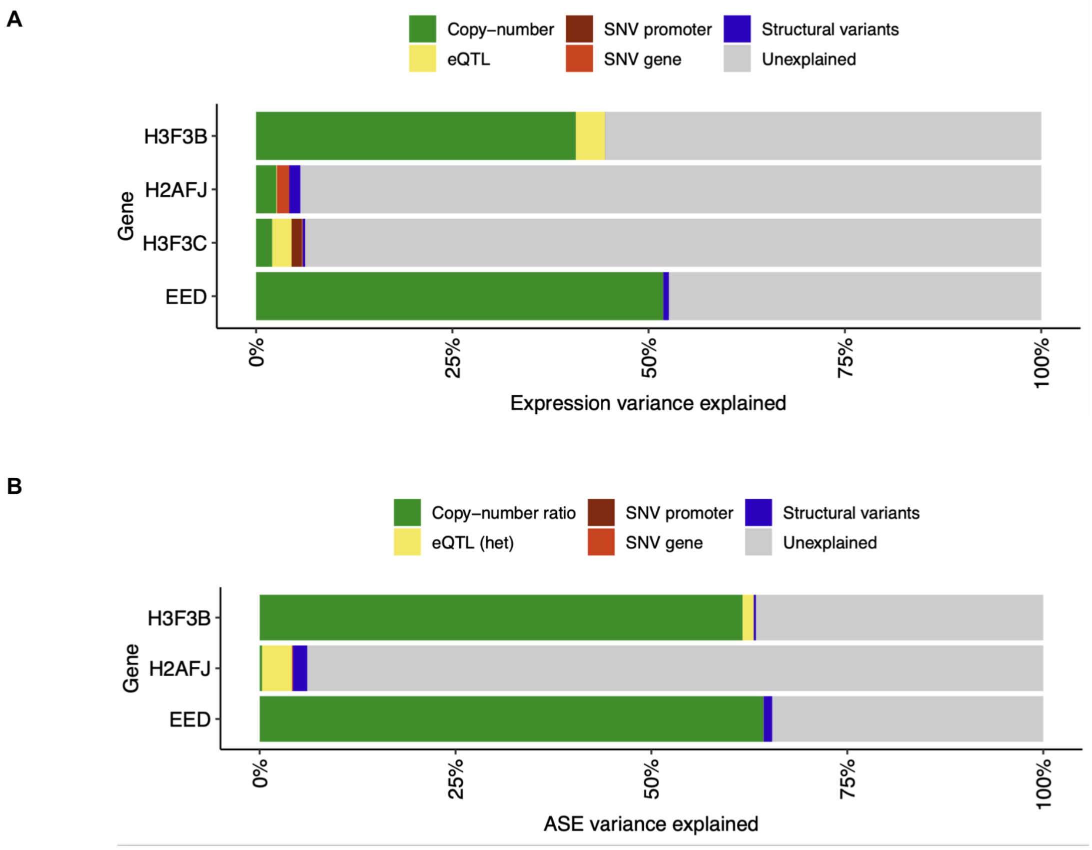


**A**, Variance in gene expression explained by genetic effects. **B**, Variance in ASE explained by genetic effects. ASE variance of gene *H3F3C* could not be investigated, because variance components were only determined for genes for which at least 20 tumors harbored at least one heterozygous expressed SNP, which is required to calculate the ASE ratio. However, heterozygous expressed SNPs within *H3F3C* were only found in a single tumor (NBL50).

#### Supplementary figure 19: H3F3B, H3F3C and H2AFJ expression in c-circle positive and negative neuroblastoma tumors

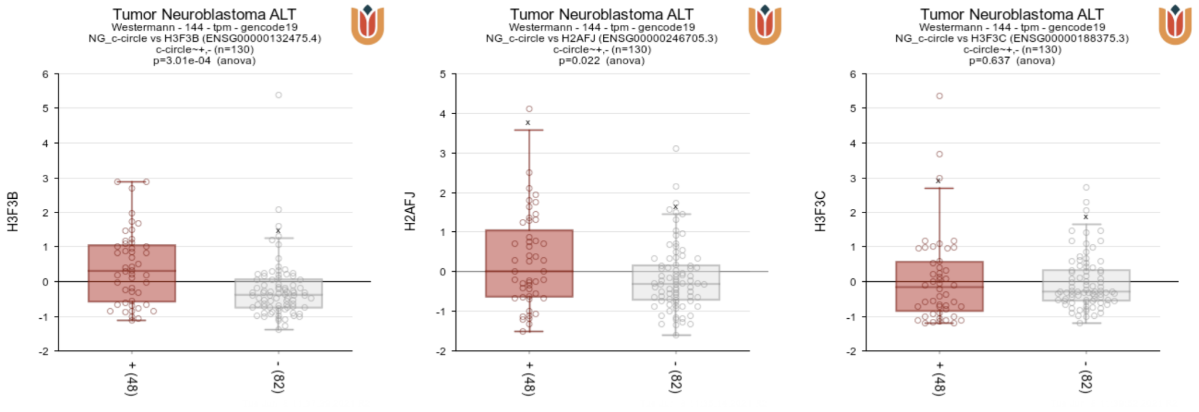


Expression z-scores of histone variant genes *H3F3B*, *H2AFJ* and *H3F3C* in 130 neuroblastoma tumors stratified by positive (+) [N=48] and negative (-) [N=82] c-circle status from Hartlieb et. al 2021 (R2: Genomics Analysis and Visualization Platform, http://r2.amc.nl).

#### Supplementary figure 20: Copy-number Log-ratio (LogR) and expression of the EED in ALT and non-ALT neuroblastomas

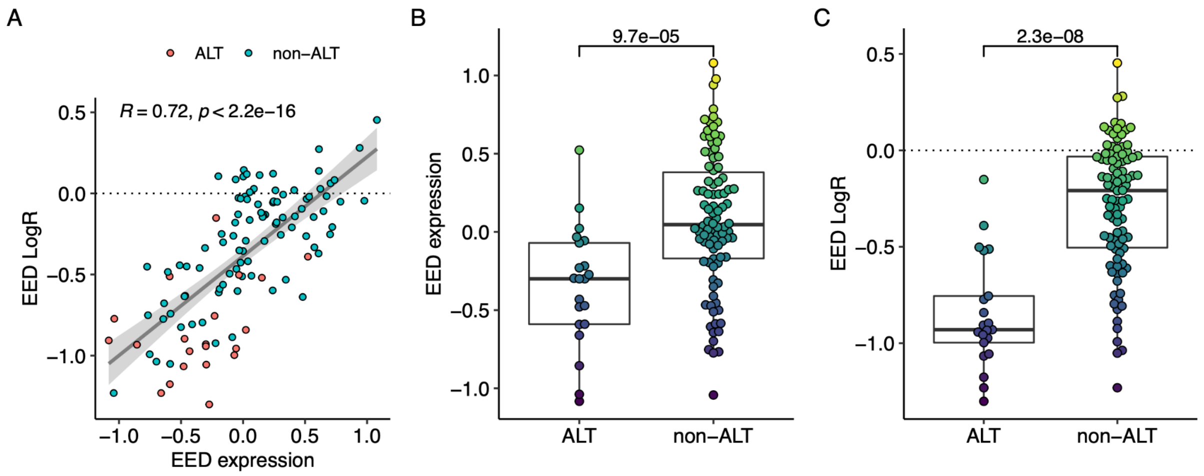


**A**, Linear regression of copy-number LogR and gene expression values for gene *EED* per tumor. Grey ribbon indicates 95% confidence interval of regression line. **B**, *EED* gene expression by ALT status. **C**, *EED* gene copy-number LogR by ALT status. P-value of Wilcox rank sum test between ALT and non-ALT tumors in (B,C) shown above bracket.

Supplementary figure 21: EZHIP expression by molecular telomere maintenance features

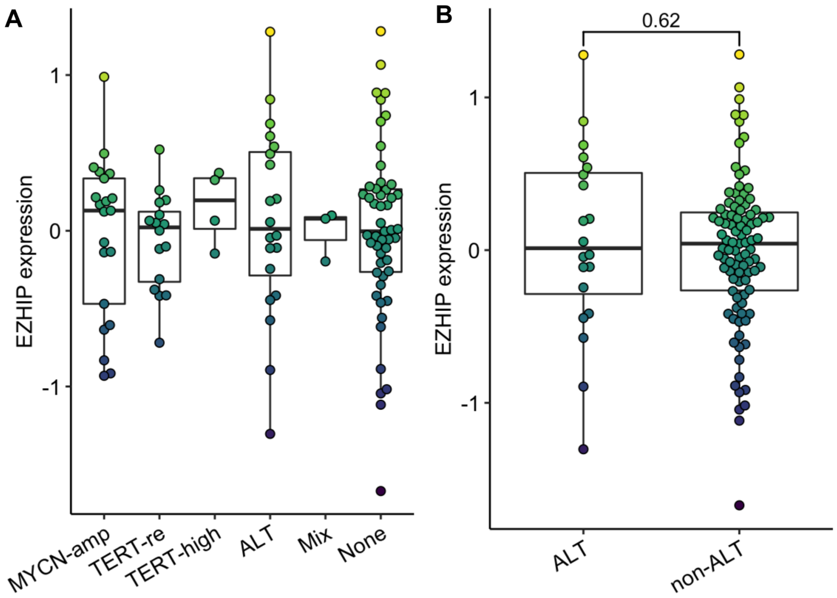


**A**, Normalized gene expression of PRC2 inhibitor *EZHIP* by molecular telomere maintenance group. **B**, Normalized gene expression of *EZHIP* by ALT status. P-value of Wilcox rank sum test between ALT and non-ALT tumors shown above bracket.

Supplementary figure 22: Contribution of local genetic effects on ASE in allelic dosage genes (top 20 genes with strongest ASE–expression correlation)

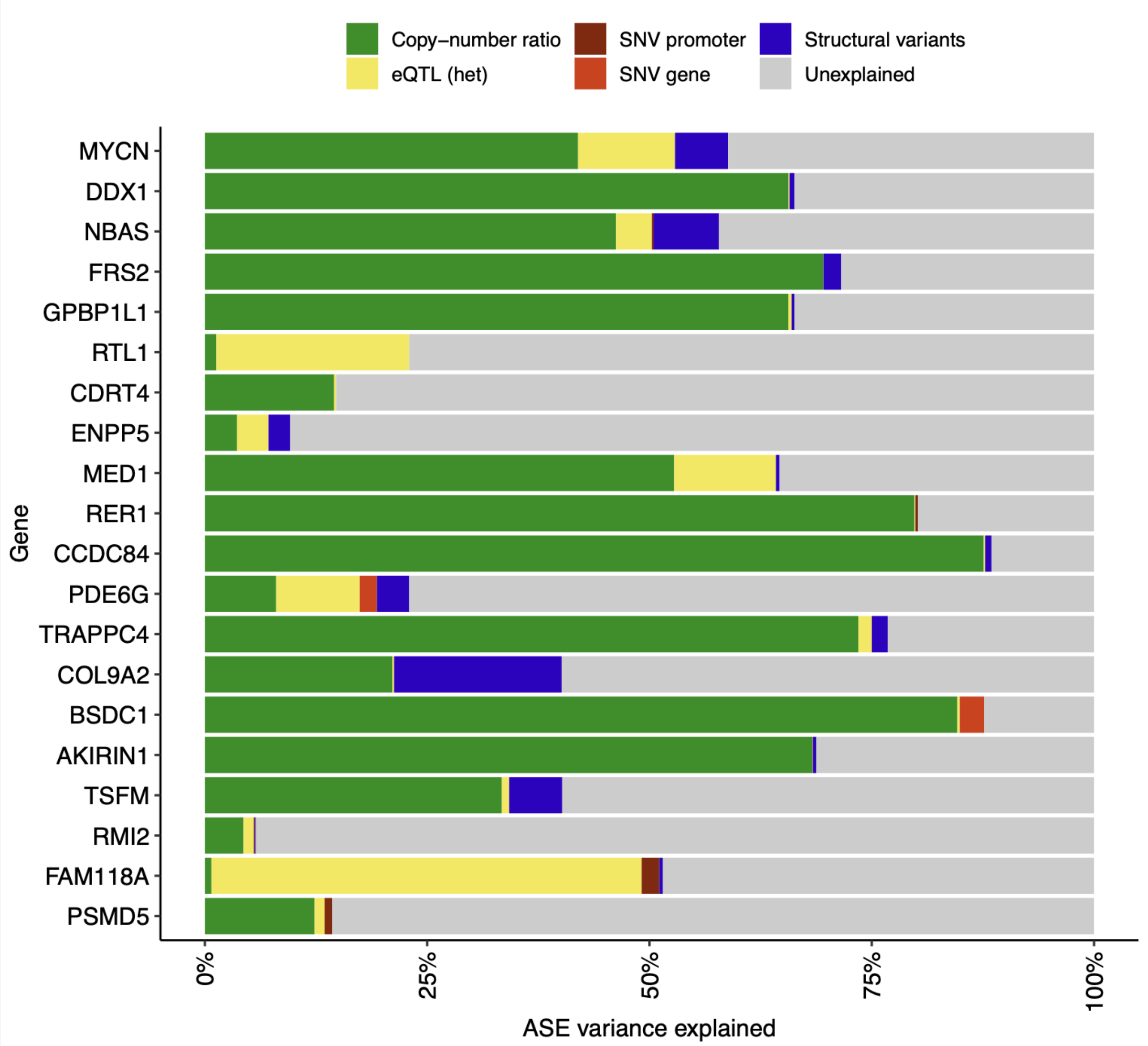

Supplementary figure 23: CpG methylation in the RTL1 upstream region

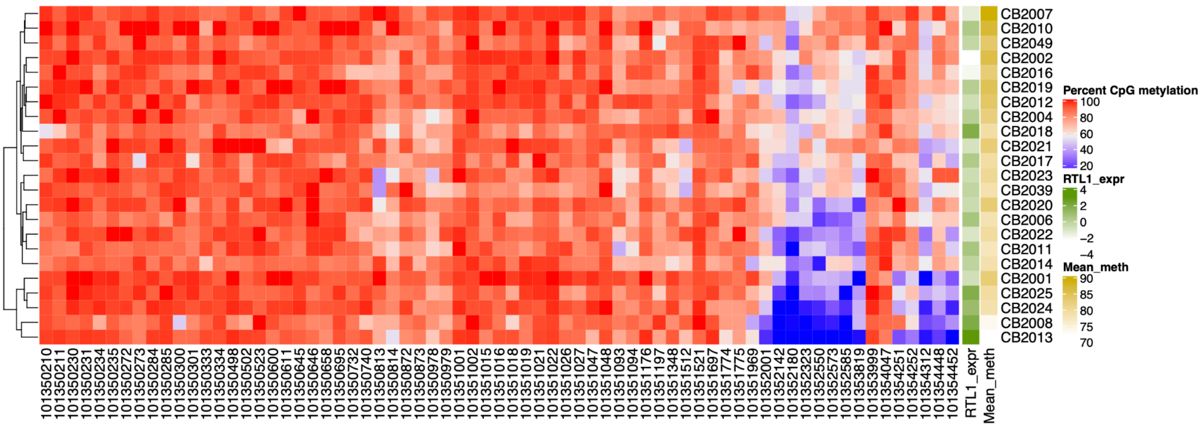


Heatmap shows percent methylated CpGs from whole-genome bisulfite sequencing of the *RTL1* upstream region in a -4kb to +1kb window relative to *RTL1* gene start. Chromosome 14 genomic location of CpG denoted on x-axis.

Supplementary figure 24: TERT promoter rearrangement in CB2064

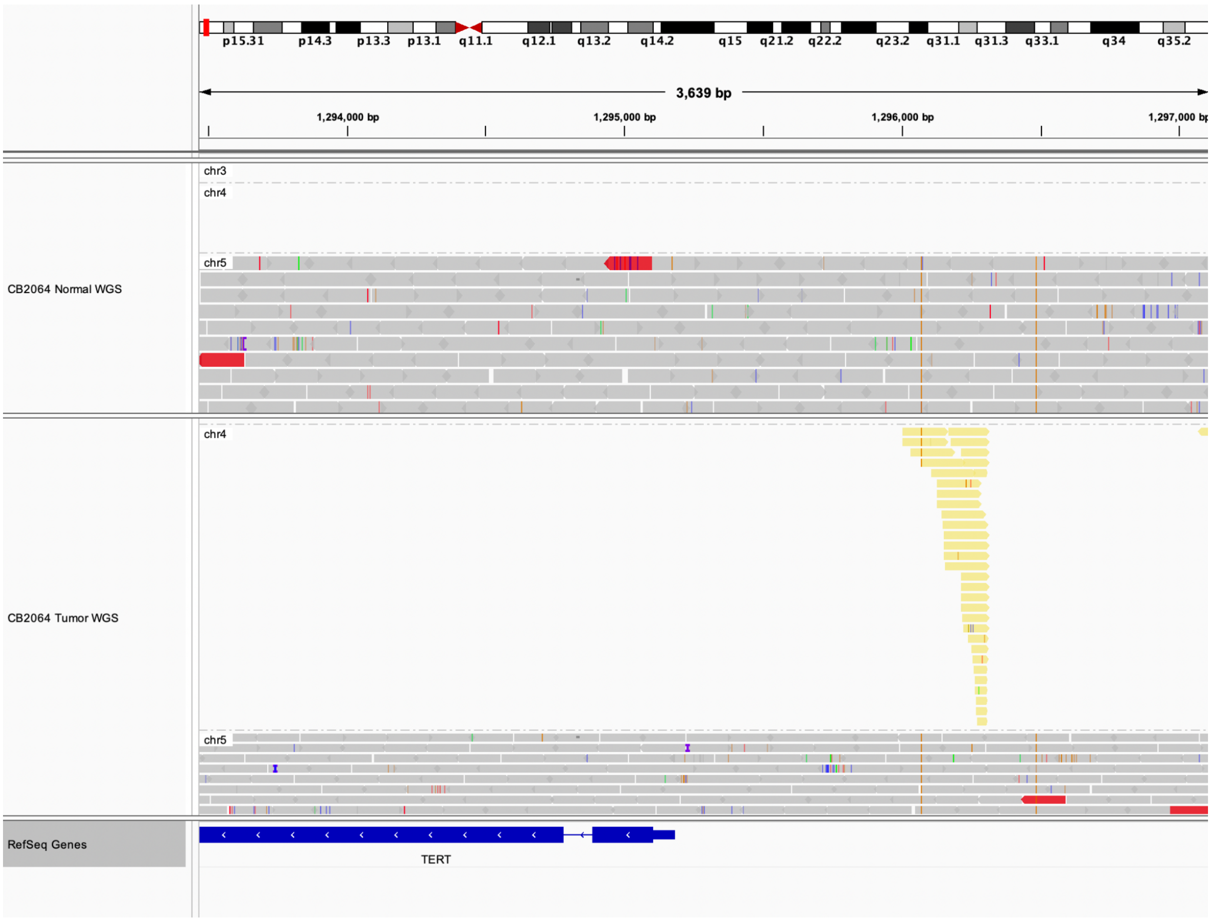


Discordant paired-end read mapping between *TERT* promoter and chromosome 4 in tumor CB2064. Aligned reads with paired-end mates mapping to chromosome 4 are indicated in yellow and found exclusively in tumor WGS (bottom) but not in normal WGS (top).

#### Supplementary tables

##### Supplementary table 1: Overview and clinical annotations of donors/samples

##### Supplementary table 2: Differentially expressed genes between deceased-from-disease and others

##### Supplementary table 3: Reactome pathways enriched in copy-number dosage effects

##### Supplementary table 4: Correlation of gene expression and 11q and 17q copy-number for differentially expressed ALT genes

##### Supplementary table 5: Allelic dosage genes

#### Supplementary methods

##### Sequencing read alignment

All reads were aligned to the GRCh37 reference genome. WGS reads were aligned with BWA-MEM 0.7.15 [2]. RNA-seq reads were aligned with STAR 2.5.3a [3]. Samblaster 0.1.24 [4] was used to mark duplicates in alignment files. Quality control was performed using FastQC. S.Table 1 lists donors from which matched tumor and normal WGS as well as matched tumor RNA-seq was used in the analyses.

##### Genotyping and phasing

Variant call files with 84,801,880 germline variants reported by the 1000 Genomes Project (phase 3)[5] were downloaded and filtered for biallelic single nucleotide polymorphisms (SNPs). SNPs from chromosome 1-22 were filtered for minor allele frequency (MAF) of 1% or higher in the 1000 Genomes cohort. A mappability signal track for 50-mers in the human reference hg19 was downloaded from the UCSC genome browser (wgEncodeCrgMapabilityAlign50mer) [6] and intersected with the SNP positions. Only SNPs with a mappability score of 1 (unique 50mer) were kept. The resulting set of 9,866,569 variant sites was defined as the *SNP panel* for further downstream analysis.

Pileups are data structures that make nucleotide base observations in sequence alignments easily accessible. For a given alignment pileups can be created for a subset of covered positions. A pileup reports the counts and identity of aligned nucleotides, mismatches and alignment gaps for each position considered. This information can be used to analyze allelic frequencies and base quality scores at predefined genomic positions, such as those of previously known SNPs. In DNA samples of healthy tissue (e.g. blood) the information of the pileup can also be used to assign genotypes to SNP positions. We generated pileups at positions of the SNP panel from whole genome sequencing (WGS) alignments of blood-derived control samples by Bcftools 1.8 mpileup, excluding unmapped reads, or reads that were marked as optical duplicates or "not primary alignment". The resulting pileups were then used as input to the Bcftools 1.8 multiallelic-caller to call genotypes at the positions of the SNP panel. Briefly, the caller determines the number of observations per allele and employs a statistical model incorporating these frequencies and read quality scores to determine genotypes (homozygous reference, homozygous alternative, heterozygous) and a genotype quality score. We only kept resulting genotypes with an allelic depth of 10 or more reads and a genotype quality of 20 or higher. The resulting individual variant files were merged and genotypes were phased by Eagle 2.4 [7] using the phased 1000 Genomes genotypes as reference. In this step each of the two alleles (reference and alternative) is assigned to an A or B allele, which represent the two parental alleles. The method is based on dependencies in allele frequencies of neighboring SNPs and makes use of pre-existing information of haplotype assignments in the reference panel. We used the phased 1000 Genomes variants as a reference panel. The resulting variant file, comprising phased genotypes of all individuals was defined as the *genotype panel* for further downstream analysis.

##### Somatic single nucleotide and structural variation calling

Somatic SNVs were called by Mutect2 version 2.2 from the GATK software package [8]. SNV calls were filtered using a panel of normals. Effects of SNVs were predicted using the Ensembl variant variation effect predictor version 101 [9] in offline mode with distance 100,000 bp. SNVs in categories missense, splice, stop, synonymous, 5' UTR and 3' UTR were summarized to gene level somatic mutation burden. Somatic SNVs annotated as promoter variants by the Ensembl variant effect predictor were considered separately. Splice, nonsense and missense variants for each gene were summarized based on the assigned consequence.

SV were called using novobreak version 1.1.3 [10] in pairs of matched tumor and normal WGS alignments. Briefly, novobreak detects SVs by analyzing k-mers that are unique to reads in the tumor sample. K-mers of tumor reads are collected and those occurring in normal samples or in the reference sequence are removed. The remaining k-mers are clustered and local assemblies of identified kmer-containing reads are generated. Consensus sequences of the assemblies are then aligned to the reference genome to infer breakpoint positions. We only kept SV calls with QUAL ≥ 30, at least 5 high quality reads in support of each breakpoint in the tumor sample, 0 reads supporting each breakpoint in the normal sample, 5 or more discordant reads per breakpoint in the tumor sample and 3 or less discordant reads per breakpoint in the normal sample. The functional effects of SVs at the *TERT* locus have been established previously [11,12] and for the detection of *TERT* SVs we relaxed the threshold on high quality reads in support of each breakpoint, requiring at least 2 of those reads to keep the SV call. Other thresholds were applied as described above. *TERT* rearrangement status was assigned to a sample positive for at least one somatic SV 100,000 kb upstream or downstream from *TERT* gene start and end coordinates (Ensembl/GRCh37) or annotated as *TERT* rearranged in [13]. Additionally, the *TERT* locus was manually examined in tumor and normal WGS alignments for structural variants that were missed by the variant calling procedure above, resulting in one additional assignment of positive *TERT* rearrangement status in tumor CB2064 (S.Fig. 24).

We used a targeted approach to identify *ATRX* exon deletions. To this end we determined read coverage at *ATRX* gene coordinates in 50 bp bins, normalized the read counts by the number of overall mapped reads and defined a tumor coverage ratio by $s_i = \log2(n_{iT}/n_{iN})$, where $n_{iT}$ and $n_{iN}$ are normalized read counts in tumor and normal for bin *i* respectively. For each matched tumor/normal pair we then fit a two-component gaussian mixture model to the signal and determine the mean and relative proportions of two hypothetical clusters, corresponding to read coverages of deleted and intact regions of the gene. Samples that harbored a signal mean difference of at least 1.5 units between the two clusters and in which the smaller cluster showed a proportion of 10% or more of the larger cluster were regarded as *ATRX* deleted. Tumors that showed either *ATRX* deletions as determined by this method, were positive for a somatic SV breakpoint inside *ATRX* gene boundaries or carried a somatic missense, nonsense or splice SNV were considered as mutant *ATRX*.

#### Estimation of whole-genome-doubling

The WGD (whole-genome-doubling) status of the samples were estimated using the phylogenetic reconstruction tool MEDICC2 [14]. For single samples MEDICC2 calculates the minimum number of evolutionary events from a diploid genotype to the copy-number states of the sample by combining loss-of-heterogeneity events, whole-genome-doubling as well as chromosome-wide and focal losses and gains. By checking whether the shortest evolutionary path from the diploid to a sample copy-number profile contains a WGD we can estimate the WGD status. As a WGD event followed by multiple losses can also be modeled by multiple gains, MEDICC2 can be conservative in its WGD estimation. We employed a bootstrap method to improve the estimation of WGD events. For this we used 100 bootstrap copy-number profiles per sample that were created by randomly drawing 22 chromosomes with replacement from the original chromosomes. If 5% or more of the bootstrap runs exhibited a WGD the sample was marked as WGD positive.

#### Classification of copy-number states

We characterized copy-number segments based on their allele-specific copy-numbers, coverage log ratio and size. Copy-number states (CN state) were assigned to each segment based on ASCAT's allele counts, ploidy estimates and LogR of coverage between tumor and normal WGS alignment. copy-number (CN) states were assigned based on conditions in the following order: *weak gain*: $CN_{total}$ > round(ploidy); *medium gain*: $CN_{total}$ > 1.5 × round(ploidy); *strong gain*: $CN_{total}$ > 2.5 × round(ploidy); *shallow loss*: $CN_{total}$ < round(ploidy); *loss*: $CN_{total}$ < 0.5 × round(ploidy). Here, $CN_{total} = CN_{major} + CN_{minor}$ , where $CN_{major}$ and $CN_{minor}$ are the allele counts of major and minor allele respectively and round(ploidy) the ploidy estimate determined by ASCAT rounded to an integer value. Copy-number state *focal amplification* was assigned to segments smaller than 10 Mb with $CN_{major} \geq 5$ and $LogR_{seg} - LogR_{contig} > 0.7$, where $LogR_{seg}$ is the mean LogR of the segment and $LogR_{contig}$ the mean LogR of the segment's chromosome (contig). Similarly we assigned a copy-number balance state (CN balance state) to each segment. For this purpose the copy-number ratio was determined as $CN_{ratio} = CN_{major} / (CN_{major} + CN_{minor})$. Then, the CN balance state *balance* was assigned if $CN_{minor} > 0$ and $CN_{major} = CN_{minor}$. CN balance state *weak imbalance* was defined as $CN_{major} > CN_{minor}$ and $CN_{ratio} \leq ⅔$, and state *strong imbalance* was defined as $CN_{major} > CN_{minor}$ and $CN_{ratio} > ⅔$. CN balance state *amplification* was defined in the same way as for the copy-number state above. Copy-number segments were marked as *LOH* if $CN_{minor} = 0$. We derived copy-number states for each chromosome arm and separately for cytoband 1p36, (which is frequenytly deleted in *MYCN*-amplified tumors) by summarizing the overlap of all segments per copy-number state in these broader regions and assigning the copy-number state of largest overlap. We applied the same procedure to assign copy-number states to genes by overlap between gene coordinates (Ensembl version 75). Gene amplification status was inferred from its copy-number state and gene-specific LogR measurements. Genes of copy-number state *focal amplification* or those with $LogR_{gene} > 2.5$ were defined as amplified, where $LogR_{gene}$ is the mean LogR across all SNPs falling within the gene's coordinates.

#### Association of copy-number ratio with survival

To associate allelic copy-number differences with survival we summarized copy-number ratio and LogR values in genomic regions. We calculated the copy-number ratio as $CN_{ratio}$ = $CN_{major}$ / ($CN_{major}$ + $CN_{minor}$), where $CN_{major}$ and $CN_{minor}$, are major and minor allele counts as determined by allele-specific copy-number analysis respectively. $CN_{ratio}$ and LogR values were summarized on the level of chromosome arms. Additionally we summarized copy-numbers in 5 Mb bins along the genome. The average value per region was defined as the mean value of CN segments overlapping the genomic region weighted by the length of overlap. 5 Mb bins overlapping focal amplifications were assigned the value of the amplified CN segment directly, dropping values of other segments overlapping the same bin. We used this strategy in order to maintain information about amplification of (focal) alterations in larger bins (e.g. 1 Mb focal amplification in 5 Mb bins).

We associated the summarized copy-number ratio per region with patient survival. For each region we tested for the association of copy-number ratio to survival using a generalized linear regression on the binary response "deceased from disease" vs. other. The test was set up to control for covariates *MYCN* amplification, age, tumor stage 4, sex, tumor purity and tumor ploidy. The association p-value was determined by an analysis of variance (ANOVA) using a Chi-Squared test. The test was carried out between a generalized linear model (GLM) of the covariates above and a second model that included the copy-number ratio in addition to these covariates. Nominal p-values determined for each region were corrected by the Bonferroni method and regions below 0.05 FWER were considered significant.

We used a Cox proportional hazard model [15] to predict overall survival from the copy-number ratio of the chromosomal region identified in the regression analysis (Methods). In contrast to the binary survival outcome, here survival times are taken into account. Subsequent significant bins in the discovery model were merged and the average copy-number ratio was determined for the merged bins by the weighted average method as described above. Survival times were predicted by the covariates copy-number ratio, *MYCN* amplification status, age, tumor stage 4, sex, tumor purity and tumor ploidy. A survival function was estimated by the Kaplan-Meier method. Here, discretized states "balance" (copy-number ratio = 0.5) and "imbalance" (copy-number ratio > 0.5) were used to split samples into two groups and to plot the corresponding survival curves.

#### Pathway enrichment of copy-number effects on gene expression

Pathway enrichment analysis of copy-number effects on gene expression was conducted using the fgsea R package [16] version 1.12.0 with inbuilt Reactome pathway definitions.

Genes were ranked based on the variance in expression explained by copy-number effects from the variance component analysis. Significant pathways were determined at FDR < 0.01. Independently enriched pathways were determined by the collapsePathways function of the fgsea R package with default parameters.

#### Differential gene expression analysis of survival status

DESeq2 1.26.0 [17] was used to perform differential expression analysis on HTseq/htseq-count 0.9.1 [18] gene counts between donors marked with survival status *deceased from disease* according to the clinical annotation file and other donors on variance stabilized raw RNA-seq counts. P-values and log-fold changes of differential expression were obtained controlling for sample covariates cohort, tumor purity, age and sex. Log-fold changes were shrunken using the apeglm method [19]. Significant genes were determined at FDR < 0.05 based on the Benjamini Hochberg-adjusted p-value from DESeq2.

#### Allele-specific expression analysis

Allele-specific RNA read counts were determined by GATK[8] (version 3.5.0) ASEReadCounter from RNA-seq alignments at heterozygous SNPs following established protocols[20]. SNPs with less than 8 total or less than 2 allelic reads were removed. Additionally, only sites that qualified as bi-allelic according to a statistical test were retained: A binomial test on the minimum allele count = min(alt, ref), number of trials (alt + ref) and hypothesized probability of success sum(non_ref_alt)/sum(raw_depth) was applied, where ref and alt are the reference and alternative allele counts, and non_ref_alt and raw_depth the non-reference/non-alternative allele count and raw read depth per site respectively. Sites for which the null hypothesis was rejected (FDR 0.05, Benjamini-Hochberg) were classified as bi-allelic. The reference allele bias was estimated by averaging over the reference allele fraction ref / (ref + alt) of all ASE sites from balanced copy-number regions per sample. We used statistical phasing information (see genotyping and phasing methods) to summarize allelic counts at exonic heterozygous SNPs of the same haplotype per gene. Only genes with a total of 10 or more counts from both haplotypes were retained. The ASE ratio for a given gene was calculated as max(A, B) / (A + B), where A and B are haplotype counts of the arbitrary A and B allele respectively. Expression imbalances per gene and sample were assessed by a two-sided binomial test using A as the number of successes, (A + B) as the number of trials and 0.5 as the hypothesized probability of success. The p-value was adjusted for multiple testing using the Benjamini-Hochberg procedure. Allelic-expression imbalance (AEI) status was assigned to observations (gene-sample pairs) for which an expression imbalance was detected at FDR 0.05.

#### Correlation analysis of allele-specific and total expression

To determine genes underlying strong cis-regulatory control by activation or attenuation of gene expression from one of the two alleles, we performed a correlation analysis between ASE and gene expression. ASE ratios were filtered, so that only ratios from 10 or more RNA-seq read counts remained. Variance stabilized RNA-seq reads were matched with ASE ratios by sample and gene. We grouped observations by gene and only considered genes with at least 10 donors informative for ASE in that gene yielding a total og 10,862 genes with sufficient number of observations. Both ASE ratio and gene expression read counts were separately corrected by batch and tumor purity by fitting linear models per gene and obtaining residuals of expression and ASE ratio that were used in the subsequent analysis. Analogous to the ASE ratio, the *tumor DNA ratio* was defined as max(A,B)/(A+B), where A and B are phased and aggregated read counts of the tumor DNA alignment of expressed heterozygous SNPs gene for the two alleles respectively. We then modeled the ASE ratio by gene expression, cohort, tumor purity, total read count at heterozygous SNP (log) and normal DNA ratio using linear regression and determined a p-value for the gene expression term by ANOVA (F-statistic). Resulting p-values were corrected using the Benjamini-Hochberg procedure. Allelic dosage genes (AD genes) were defined as those genes at FDR < 0.05. To identify a subset of AD genes with clinical relevance we matched Pearson's r of ASE-expression correlation with adjusted P values from differential expression analysis. A subset of differentially expressed genes was defined by intersecting AD genes with genes significantly different expressed between deceased and not-deceased patients (FDR < 0.05, Benjamini-Hochberg).

#### cis-QTL association testing

For eQTL analysis the SNP genotypes called in 115 WGS samples of normal tissue were pooled and filtered. Only SNPs with a minor allele frequency of 5% and at least 10% genotyped samples in the cohort were retained. Htseq count [18] was used to count reads from RNA-seq data of tumor samples in the union of all exons per gene based on the Ensembl 75 gene annotation. Raw RNA gene counts were normalized by library depth per sample and transformed to variance-stabilized counts by DESeq2 [17]. Only protein-coding genes on chromosomes 1-22 with at least 10 counts in 90% of the samples were considered. In total 13,903 genes were included in the analysis. Variance-stabilized counts were centered and strong outlier samples, defined as normalized count values exceeding 3 times the standard deviation of all normalized counts per gene, were removed. To estimate the expression variability between samples we applied probabilistic estimation of expression residuals (PEER) [21] to derive 10 factors from the normalized counts. We took these factors as representatives for global expression differences, that are likely not associated with cis-regulatory effects and incorporated them as covariates in the association test described below. Genotypes of SNPs in a cis-window of 500kb upstream and downstream of annotated gene coordinates were associated with the gene's quantiative trait. SNPs were associated with quantitative traits by FastLMM [22], version 0.2.23 in single SNP mode. FastLMM uses a linear mixed model in a regression of the number of alternative alleles on quantitative phenotypes controlling for given covariates. We combined gene- and sample-specific covariates individually in each test. The somatic gene copy-number was calculated as the average total copy-number in gene intervals and used as the only gene-specific covariate. Sex, cohort, tumor purity, tumor ploidy and the 10 PEER factors were incorporated as sample-specific covariates. Each association test was controlled by the matching set of sample and sample-gene-specific covariates for a given set of gene associations.

#### Protein network visualization

Protein networks visualizations were created using the STRING DB network viewer (version 11.0b, https://version-11-0b.string-db.org/) [23]. Settings were adjusted such that line thickness between nodes indicate protein interaction confidence (strength of data support) and minimum required interaction score was set to "medium" (0.4). The default set of interaction sources was used (text mining, experiments, databases, coexpression, neighborhood, gene fusion, cooccurrence).

The interaction graph in ALT associated protein interactions of ATRX and histone variant genes was subset to proteins of differentially expressed genes (ALT) (FDR < 0.05) with correlation to LogR of 11q or 17q with abs(*r*) > 0.3, where *r* is Pearson's correlation coefficient. Network nodes were colored based on up- (red) or down- (blue) regulated genes as identified in ALT differential expression analysis. Enrichment of biological processes (GO terms) in the protein network of 17p dosage effects were determined by the STRING network analysis tool.

#### Whole-genome bisulfite sequencing analysis

Whole-genome bisulfite sequencing data was obtained for a subset of 23 tumors (S.Table 1). Libraries were prepared using the EpiTect Bisulfite and Illumina Truseq PCRFree DNA sequencing kit (V2.5) and sequenced on the Illumina Hiseq X plattform yielding paired-end reads of 2×150 bp. Reads were extracted with bcl2fastq (v.2.19.0.316) and processed with a developer version of the PiGx BS-seq pipeline [24]. Quality control was performed with FastQC 0.11.9. Reads were trimmed with TrimGalore v.0.6.2 and aligned to the bisulfite-converted reference human genome GRCh37 using bwa-meth v.0.7.17 [25]. Duplicate reads were removed with samblaster v.0.1.24 [4]. Germline C/T SNPs detected in our cohort at MAF > 0.01 were removed using bcftools 1.9. CpG DNA methylation calling was performed with methylKit v.1.15.4 [26] at a minimum 10x coverage.
